# Whole-genome evaluation of genetic rescue: the case of a curiously isolated and endangered butterfly

**DOI:** 10.1101/2024.12.20.627963

**Authors:** Zachary G. MacDonald, Julian R. Dupuis, James R. N. Glasier, Robert Sissons, Axel Moehrenschlager, H. Bradley Shaffer, Felix A. H. Sperling

## Abstract

Genetic rescue, or the translocation of individuals among populations to augment gene flow, can help ameliorate inbreeding depression and loss of adaptive potential in small and isolated populations. Genetic rescue is currently under consideration for an endangered butterfly in Canada, the half-moon hairstreak (*Satyrium semiluna*). A small, unique population persists in Waterton Lakes National Park, Alberta, isolated from other populations by >350km. However, whether genetic rescue would actually be helpful has not been evaluated. Here, we generate the first chromosome-level genome assembly and whole-genome resequence data for the species. We find that the Alberta population’s genetic diversity is extremely low and very divergent from the nearest populations in British Columbia and Montana. Runs of homozygosity suggest this is due to a long history of inbreeding, and coalescent analyses show that the population has been small and isolated, yet stable for up to 40k years. When a population maintains its viability despite inbreeding and low genetic diversity, it has likely undergone purging of deleterious recessive alleles and could be threatened by their reintroduction via genetic rescue. Ecological niche modelling indicates that the Alberta population also exhibits environmental associations that are atypical of the species. Together, these results suggest that population crosses are likely to result in outbreeding depression. We infer that, in this case, genetic rescue has a relatively unique potential to be harmful rather than helpful at present. However, due to reduced adaptive potential, the Alberta may still benefit from future genetic rescue as climate conditions change. Proactive experimental population crosses should be completed to assess reproductive compatibility and offspring fitness.

## Introduction

Quantifying whole-genome genetic diversity and patterns of differentiation within endangered species is an integral part of modern conservation practice (Kardos et al. 2021). This is particularly true for small and isolated populations, which are often characterized by high inbreeding, low genetic diversity, and low adaptive potential (Lande 1988; Saccheri et al. 1998; Kyriazis et al. 2023). The evolutionary dynamics of these populations are of ever-increasing relevance to conservation, particularly as anthropogenic habitat fragmentation continues to break apart naturally connected populations (Haddad et al. 2015; Schlaepfer et al. 2018; MacDonald et al. 2021; 2024). It is often uncertain whether small, isolated, but ecologically important populations will persist without intervention, or if deleterious genetic processes, primarily inbreeding depression, will lead to their extinction (Keller and Waller 2002; Charlesworth and Willis 2009; Hedrick and García-Dorado 2016; Whitla et al. 2023).

Inbreeding is a significant threat to small populations due mainly to the stochastic homozygous expression of deleterious recessive alleles, although loss of heterozygote advantage and epistasis can also be important (Charlesworth and Willis 2009; Hedrick and García-Dorado 2016). Following Kyriazis et al. (2023), we define “inbreeding load” as the quantity of deleterious recessive alleles in an individual or population that are masked in heterozygous form (Hedrick and García-Dorado 2016) and “genetic load” as the realized reduction in fitness when these alleles are expressed in homozygous form (Kirkpatrick and Jarne 2000; Bertorelle et al. 2022). If an effective population size is sufficiently small, random mating among related individuals can quickly convert inbreeding load into genetic load and become a primary determinant of extinction risk (Kyriazis et al. 2021). Additionally, drift may overwhelm purifying selection against weakly deleterious alleles (i.e., with selection coefficients <<1 / (2*N_e_*)), allowing them to become fixed (Kimura et al. 1963; Lynch et al. 1995; Robinson et al. 2022). This “fixed load” results in permanent reductions in fitness unless new alleles are introduced (Charlesworth 2018). A viable conservation practice used to reduce inbreeding depression and fixed load is genetic rescue, which aims to increase genetic diversity by (re-)establishing gene flow between isolated populations (Storfer 1999; Weeks et al. 2011; Ralls et al. 2020). Genetic rescue and related forms of population crosses have effectively alleviated inbreeding depression in a number of well-known taxa, such as the Scandinavian wolf (*Canis lupus*; Vila 2003), Florida panther (*Puma concolor coryi*; Pimm et al. 2006), greater prairie chicken (*Tympanuchus cupido*; Mussmann et al. 2017), African lion (*Panthera leo*; Trinkel et al. 2008), Glanville fritillary butterfly (*Melitea cinxia*; Mattila et al. 2012), and jellyfish tree (*Medusagyne oppositifolia*; Finger et al. 2011) (and see review by Clarke et al. 2024). Theoretically, genetic rescue can also increase the standing genetic diversity of a population, increasing its adaptive potential to survive future environmental change (Willi et al. 2006; Mable 2019). Although real-world examples of genetic rescue actually increasing the adaptive potential of wild populations are lacking, it remains an important consideration and motivation for translocations.

Despite clear benefits, translocations also entail the theoretical risk of introducing new inbreeding/genetic load into already threatened populations (Hedrick et al. 2014; 2019). In small and isolated populations, a principal consequence of inbreeding is the reduced fitness and demographic loss of individuals due to the homozygous expression of deleterious recessive alleles. However, the reduced fitness or death of these individuals also reduce the frequency of those recessive alleles and thus the consequences of future inbreeding—a process known as genetic purging (Glémin 2003; Xue et al. 2015; Hedrick and Garcia-Dorado 2016; Robinson et al. 2016; 2018; Grossen et al. 2020; López-Cortegano et al. 2021; Pérez-Pereira et al. 2021; Mooney et al. 2023; Pečnerová et al. 2024). If purging has occurred, genetic rescue may theoretically reintroduce inbreeding load to the detriment of the recipient population. Theory suggests that small to moderate population sizes are optimal for purging, wherein gradual, continuous inbreeding is sufficient to consistently expose genetic load to purifying selection without compromising population viability (Day et al. 2003; Glémin 2003; García-Dorado 2012; Pekkala et al. 2012; Robinson et al. 2016; 2018; Kyriazis et al. 2021; Pérez-Pereira et al. 2021). However, what constitutes an optimal population size depends on both the genetic diversity and demographic history of a population (Caballero et al. 2017; Mable 2019; Robinson et al. 2022). Adding further confusion, inbreeding load, genetic load, and the process of purging have proven extremely difficult to quantify in nature, most often relying on simulations with nebulous interpretations (Leber and Firmin 2008; Frankham et al. 2020; Kyriazis et al. 2023). Notwithstanding, it remains the case that heathy, resilient populations with a long history of complete isolation, high homozygosity, and no signs of inbreeding depression have likely purged much of their genetic load (Robinson et al. 2016; 2018; 2022; Pečnerová et al. 2024). Although these populations may also harbor a fixed load, their demographic health and resiliency indicate that any associated fitness consequences are overtly insufficient to compromise population viability. From this perspective, small populations with very long histories of isolation may not immediately benefit from, and may even be harmed by, genetic rescue (Hedrick et al. 2014; Robinson et al. 2016; 2018; Hedrick et al. 2019; López-Cortegano et al. 2021; Kyriazis et al. 2021; 2023; Mooney et al. 2023).

Another risk associated with genetic rescue is outbreeding depression, defined as reductions in the fitness of translocated individuals’ progeny due to the inheritance of maladapted phenotypes or the disruption of favourable gene combinations (Templeton 1986; Tallmon et al. 2004; Edmands 2007; McBride and Singer, 2010). According to Frankham et al. (2011), the probability of outbreeding depression is substantial when donor and recipient populations exhibit at least one of the following attributes: 1) they are different species; 2) they exhibit fixed chromosomal differences; 3) they have experienced no gene flow over the last 500 years; or 4) they inhabit different environments. In wild populations, attribute four—inhabiting different environments—is particularly important because it implies divergent adaptation to different environmental/ecological conditions (Nosil 2012; MacDonald et al. 2020; Campbell et al. 2022; Grether et al. 2024). Even if populations show little neutral genetic differentiation, moving individuals across gradients of selection can result in a substantial loss of fitness, as has been shown in Edith’s Checkerspot butterfly (*Euphydryas editha*; Singer and McBride 2010; McBride and Singer 2010; review in Parmesan et al. 2023). However, even though crosses among divergent populations result in overall genomic homogenization, important alleles associated with local adaptation may not always be eliminated by gene flow (e.g., Fitzpatrick et al. 2020). Further, there are counter-cases wherein crossing divergent populations results in increased fitness, a phenomenon known as both heterosis and hybrid vigor (Darwin 1859; Birchler et al. 2003; Lippman and Zamir 2007). Clearly, there is a need to evaluate the risks of outbreeding depression on a case-by-case basis, carefully considering the ecology and evolution of the populations in question.

In this study, we applied whole-genome resequencing and ecological niche modelling to an endangered butterfly, the half-moon hairstreak (*Satyrium semiluna* Klots, 1930, family: Lycaenidae). Our goals were to quantify patterns of genetic diversity and differentiation within the species at its northern range limit, as well as to infer the demographic history and assess niche divergence of a small, isolated population of critical conservation concern. The range of *S. semiluna* spans the southern interior of British Columbia and southwestern Alberta, Canada, south to the eastern slopes of the Californian Sierra Nevada, and east to eastern Wyoming and northern Texas in the USA. While the species is “apparently secure” across its USA and global range (COSEWIC 2006; ECCC 2016; NatureServe 2024), less than 1% of the species’ range is in Canada, where it is listed as federally endangered. All but one of the Canadian populations occur in south central British Columbia, with an aggregate abundance qualitatively estimated to be between 5,000 and 15,000 individuals (COSEWIC, 2006). These populations, as well as those throughout the USA, inhabit steppe-like habitats dominated by big sagebrush (*Artemisia tridentata*). Approximately four-hundred kilometers to the east, there exists a single *S. semiluna* population that persists on a ∼300 ha alluvial fan (Blakiston Fan) in Waterton Lakes National Park, Alberta (COSWEIC 2006). The habitat occupied by this population is unique for the species, described as prairie/grassland dominated by sedges, grasses, and herbaceous plant species with few shrubs. Larval host plant associations also vary. British Columbia and many Pacific Northwest populations have been observed to feed on silky lupine (*Lupinus sericeus*), whereas the Alberta population feeds only on silvery lupine (*Lupinus argenteus*), despite silky lupine being readily available at Blakiston Fan.

The historical size of the Alberta population is unknown, but qualitatively estimated to have been between 2,000 and 10,000 individuals (COSWEIC 2006). Historical surveys suggest that this population fluctuates in density, with rough estimates ranging from 20.1 individuals/ha in 2004 (Kondla 2004; COSEWIC 2006) to 62.9 individuals/ha in 2008 (unpublished data). In 2017, a precipitous decline was recorded, with only 1.5 individuals/ha observed. This decline continued in 2018 with 0.5 individuals/ha, following an intense wildfire in the fall of 2017 (Eisenberg et al. 2019) that burned approximately 50% of *S. semiluna* habitat. While the cause of this decline is not known, the population has since recovered, with density estimates for 2021, 2022, and 2023 of 30.6, 8.1 and 7.9 individuals/ha, respectively (unpublished data). In truth, lack of standardized survey methods and effort across years renders estimates of population size and trends approximate at best. However, experts still generally agree that the Alberta population is small, likely completely isolated, and unique in its habitat associations (Environment and Climate Change Canada 2016).

Translocation of individuals from the nearest populations in either British Columbia or Montana is currently being considered by Parks Canada as an option to bolster genetic diversity, reduce potential inbreeding, and increase the population size at Blakiston Fan. To better inform whether this conservation intervention is warranted, we generated a chromosome-level reference genome for *S. semiluna*, representing the first high-quality assembly for the subfamily Theclinae, and whole-genome resequencing data for 15 individuals collected from Alberta, British Columbia, and Montana populations. These data were used to quantify patterns of genomic diversity and differentiation within and among populations, as well as infer each population’s inbreeding and demographic history. We also used ecological niche modelling to quantify niche divergence within the species, assessing whether population crosses are likely to result in outbreeding depression. Together, these analyses demonstrate how a substantial amount of information can be gleaned from whole-genome resequencing of a very small sample of individuals—a critical consideration when working with endangered species—and the utility of integrating genomic data with ecological modelling to inform genetic rescue decisions.

## Materials and Methods

### Sampling

We collected a total of 19 adult *S. semiluna* using aerial nets throughout the summer of 2021, preferentially collecting worn individuals when possible to minimize impacts on populations. Eight individuals were collected from Blakiston Fan, Waterton Lakes National Park, Alberta, Canada, four from Richter Pass, British Columbia, Canada, three from Anarchist Mountain, British Columbia, and four near Red Lodge, Montana, USA. Specimen metadata are available in Supporting Information Table S1. The two British Columbia collection locations were separated by approximately 17 km and individuals from both are treated here as a single population. All specimen metadata are reported in Supporting Information. Specimens from Waterton Lakes National Park were collected under the Parks Canada Agency Research and Collection Permit: WL-2021-39020. Specimens collected in British Columbia were collected on private land with landowner permissions, Nature Conservancy Canada Research Permit No. NCC_BC_2021_SS001, and Nature Trust of British Columbia #3461.

### Reference genome sequencing and assembly

Four of the eight Alberta individuals were used to generate a chromosome-level reference genome for *S. semiluna*. Genome sequencing and assembly was accomplished using a combination of PacBio HiFi long-read sequencing (Pacific BioSciences, Menlo Park, California, USA) and Omni-C proximity ligation (Dovetail Genomics, Scotts Valley, California, USA). We opted to build a reference genome based on the homogametic sex to maximize the probability of successfully sequencing and assembling the Z chromosome. Detailed methods for sequencing and assembly are provided in Supporting Information and data are available under NCBI BioProject PRJNA907836. Default parameters/settings were used for all analyses unless specified.

To assess completeness of our reference genome, we ran BUSCO v5.2.2 (Manni et al. 2021) using default parameters and the lepidoptera_odb10 lineage dataset, composed of 5,286 genes. We also quantified and classified repetitive content, including transposable elements, endogenous retroviruses, and repeat motifs, using RepeatMasker v4.1.4 (Smit et al. 2015). A library of repetitive elements for Lepidoptera was compiled using Dfam v3.6 (Hubley et al. 2016). HMMER (nhmmscan v3.3.2; hmmer.org) was used to search the reference sequence against this library using default settings, including categorization of simple repeats. Using the RepeatMasker General Feature Finding (-gff) output, we also constructed a bed file (see Supporting Information) that identifies the locations of all repetitive elements. Methods for identifying the Z chromosome in our assembly using BUSCO genes are available in Supporting Information.

### Whole-genome resequencing

Short-read, whole-genome resequencing was completed on 15 individuals. Preservation methods are reported in Supporting Information Table S1. We extracted genomic DNA from thoracic tissue using DNeasy Kits (Qiagen, Hilden, Germany), following the manufacturer’s protocol with the addition of a bovine pancreatic ribonuclease A treatment (RNaseA, 4 μl at 100 mg/ml; Sigma-Aldrich Canada Co., Canada). Following extraction, genomic DNA was ethanol precipitated and stored in purified (50 μl Millipore) water at −20°C. PCR-free whole-genome library preparation was completed using an Ultra II FS DNA Library Prep Kit (New England Biolabs, Ipswich, MA, USA), followed by paired-end, 150-bp sequencing on an Illumina NovaSeq S1 300 flowcell (total output 1600M read pairs, 500Gbp), aiming for ∼20x coverage per sample, at the Centre for Health Genomics and Informatics, University of Calgary.

### Short read processing, alignment, and genotyping

We processed raw reads using a pipeline based on recommendations from Genome Analysis Toolkit (GATK) Best Practices Guides (Van der Auwera and O’Connor 2020). After trimming adapter sequences and individual indexes, we aligned reads to our *S. semiluna* reference genome using BWA-MEM2 (Vasimuddin et al. 2019). Following removal of duplicate reads in BAM files using MarkDuplicates (Picard), alignments were passed to GATK’s HaplotypeCaller, which assembles local de-novo haplotypes on an individual-by-individual basis, generating an intermediate GVCF file for each individual. GVCF files were then used in GATK’s GenotypeGVCFs for joint genotyping of all individuals, using the “-all-sites” option. From the resulting multisample VCF file, we removed loci occurring on small, unassembled scaffolds (<1 Mb) and the Z chromosome, leaving only loci that occur on assembled autosomes. Filtering was completed using VCFtools 0.1.14 (Danecek et al. 2011), including the removal of indels, sites with >2 alleles, and sites with less than 99.9% accuracy (Phred scores < 30). We then applied an individual-specific read depth filter, removing loci with depths less than five or exceeding the 99^th^ percentile of each individual. Finally, we removed loci with more than 25% missing data across all individuals. Unless otherwise specified, a final VCF file including only variant sites (single nucleotide polymorphisms; SNPs) was used in subsequent analyses.

### Genetic divergence and population structure

To assess genetic divergence among the three collection locations, we first estimated Weir and Cockerham’s *F*_ST_ (Weir and Cockerham, 1984) using VCFtools. We also estimated *d_xy_*, the average number of nucleotide differences between individuals from two different populations (Nei and Li, 1979), using pixy (Korunes and Samuk 2021). Estimation of *d_xy_* was completed using a window size of 10 KB and included both variant and invariant sites to avoid bias caused by interpopulation variation in the amount of missing data (Korunes and Samuk 2021). Following pixy recommendations, average *d_xy_* was estimated by summing raw counts and recomputing differences/comparisons ratios, rather than averaging summary statistics across windows.

Population genetic structure was further assessed using a combination of principal component analysis (PCA) on genomic data using the R package adegenet v.2.1.1 (Jombart, 2008) and the model-based clustering program structure 2.3.4 (Pritchard et al. 2000). For both analyses, we used a sliding window to thin SNPs to a maximum density of one per 10 KB to minimize physical linkage, which has been documented to decay to baseline within 1 – 10 KB in *Heliconius* butterflies (Martin et al. 2013) and near-baseline in 100 bp in the Monarch butterfly (*Danaus plexippus*; Zhan et al. 2014). For structure analyses, ten independent runs were completed for each value of *K* = 1 – 4 using the admixture model and correlated allele frequencies. We used standard settings: the burn-in period and number of Markov chain Monte Carlo (MCMC) repetitions were set to 100,000 and 1,000,000, respectively, location priors were set to collection localities (n = 4) to inform the MCMC algorithm without biasing the model, and the alpha prior (relative admixture levels between populations) was set to one divided by the number of assumed populations (Alberta, British Columbia, and Montana). We also ran structure hierarchically, taking discrete clusters identified at *K* = 2 (membership based on admixture proportion >0.8) and rerunning them independently. In all runs the optimal value of *K* was inferred using both the Δ*K* method (Evanno et al. 2005) and the rate of change in the likelihood of *K* from 1:4 (Pritchard et al. 2000).

### Genetic diversity and runs of homozygosity

Following Kyriazis et al. (2023), we estimated heterozygosity as the proportion of heterozygous sites for each individual in nonoverlapping 1 MB windows across the autosomal genome, allowing visualization of variation in heterozygosity both within and among individuals. We also estimated population π, the average number of nucleotide differences between individuals in the same population (Nei and Li, 1979) using pixy.

We quantified runs of homozygosity (ROH) for each individual using BCFtools/RoH (Narasimhan et al. 2016), relying on genotype calls (-G30 flag) and alternate allele frequencies estimated using all individuals. The proportion of an individual’s genome contained within ROH (*F*_ROH_) is often used as a relative estimate of the prevalence of inbreeding. When millions of SNPs are analyzed (as is the case here), short ROH can be identified, meaning *F*_ROH_ can serve as both a measure of both historical and recent inbreeding, with the added benefit distinguishing between the two based on the distribution and abundance of run lengths (Kardos et al. 2017; Kardos et al. 2018b). ROH calls were categorized as short (0.1 – <0.25 MB), medium (0.25 – <0.5 MB), long (0.5 – <1 MB), or very long (>1 MB). *F*_ROH_ was estimated as the total length of all ROH calls >0.1 MB divided by 1,187,570,293, the cumulative length of autosomes within our *S. semiluna* reference genome. In theory, this statistic approximates the proportion of the genome for which both copies of the region are identical by descent, a product of either recent inbreeding (long ROH) or historical inbreeding (short ROH).

We also used VCFtools to estimate an inbreeding coefficient as *F* = (0−E)/(N−E), where *O* is the observed number of homozygous SNPs in a sample, *E* is the expected number of homozygous SNPs based on allele frequencies across all individuals and Hardy-Weinberg equilibrium, and *N* is the total number of SNPs per individual. This is effectively a measure of homozygosity relative to Hardy–Weinberg expectations and referred to as *F*_HOM_. In contrast to *F*_ROH_, which is bounded between 0 and 1, *F*_HOM_ exhibits higher variance and can take on negative values, which indicate excess heterozygosity relative to Hardy–Weinberg expectations (Kardos et al. 2018a).

### Demographic history and effective population size

Historical fluctuations in *N*_e_ were estimated from 2.5 mya until 5 kya using Pairwise Sequentially Markovian Coalescent (PSMC) analyses (Li and Durbin 2011). PSMC relies on coalescence-based expectations to estimate demographic history from single, unphased diploid genomes. Historical fluctuations in *N*_e_ for the Alberta, British Columbia, and Montana populations were estimated using each of the 15 individuals independently. We set the generation time to 1□year, as *S. semiluna* is known to be univoltine, and the mutation rate to 2.9□×□10-9 mutations per year, based on the best estimate for *Heliconius melpomene* (Keightley et al. 2015). Methods used to generate a consensus sequence for each individual and parameters used to run PSMC are available in Supporting Information.

PSMC cannot estimate contemporary *N*_e_ or recent demographic changes, e.g., within the last 10,000 years in humans (Li and Durbin 2011). To estimate contemporary *N*_e_, we used currentNe (Santiago et al. 2023), which relies on linkage disequilibrium among SNPs for multiple individuals sampled from a single population. Using both simulations and comparisons to other methods, Santiago et al. (2023) demonstrate that this method provides reliable estimates and confidence intervals, even when sample sizes are small. If the locations of SNPs are known, currentNe provides *N*_e_ estimates based on LD between SNP pairs located on different chromosomes for maximum accuracy. Inputs for currentNe were generated by subsetting our VCF file by location (Alberta [*n* = 4], British Columbia–Richter Pass [*n* = 4], British Columbia–Anarchist Mountain [*n* = 3], and Montana [*n* = 4]), and then randomly selecting ∼2 million biallelic SNPs with no missing data (the maximum number of SNPs permitted by the software).

### Niche analyses and prediction of suitable habitat

To quantify the ecological and environmental niche of *S. semiluna* and predict suitable habitat across the central and northern extent of the species’ range, we parameterized ecological niche models using MaxEnt software (Phillips et al. 2006) implemented via the R package dismo v1.3-3 (Hijmans et al. 2011). This approach uses machine-learning maximum entropy modelling to quantify ecological and environmental associations using occurrence (presence-only) records and geographic information systems (GIS) predictor variables. We generated a rectangle around the collection locations of our sequenced individuals buffered by 200 km to define the study area (Phillips and Dudík 2008; Fourcade et al. 2014), encompassing much of the central and northern extent of the species’ range. Occurrence records used for model parametrization included the collection locations of sequenced individuals (*n* = 4) and research-grade *S. semiluna* iNaturalist observations within the study area (duplicate locations removed, *n* = 74), downloaded via the Global Biodiversity Information Facility (GBIF.org accessed 17 November 2023). We used iNaturalist occurrences due to the relatively high reliability of research-grade species identification, consistency in geographic accuracy (filtered for a maximum inaccuracy of 100 meters), and large number of recent records (filtered for 2015 or later) (Beninde et al. 2023). Following MacDonald et al. (2022), geographical predictor variables included terrain ruggedness, heat load (based on terrain slope and aspect), and land cover (12 categories). Terrain ruggedness and heat load indices were estimated using the R packages raster (Hijmans and van Etten 2016) and spatialEco (Evans 2021), respectively, using a digital elevation model (Wang et al. 2016). Land cover GIS data were acquired from the Commission for Environmental Cooperation (http://www.cec.org/), generated using 2015 Landsat satellite imagery. We compiled 17 environmental variables of interest using ClimateNA v7.30 software (Wang et al. 2016; downloaded via AdaptWest Project, 2022; see Supporting Information for variable list). We did not remove correlated variables or perform variable reduction via principal component analysis (PCA; e.g., in MacDonald et al. 2022), given that MaxEnt model training is robust to predictor collinearity and accounts for redundant variables (Feng et al. 2019). Collinearity has been shown to be problematic only in terms of model transfer, e.g., in making predictions across space or time to different environmental conditions (Guisan and Thuiller 2005; Elith and Leathwick 2009; Peterson et al. 2011). All GIS data layers were reprojected to an equal-area projection (Lambert Conformal Conic) at 1 km resolution, matching the native projection of the AdaptWest dataset. Five different MaxEnt models were run, each time withholding a different 20% of occurrences for cross-validation and receiver operating characteristic (ROC).

To assess niche divergence of the Alberta population relative to other populations throughout the central and northern extent of the species’ range, we parameterized another MaxEnt model using the same methods, but this time removing all occurrences representing the Alberta population. This model was then used to predict habitat suitability at the locations of all occurrences including the Alberta population. If the predicted suitability value of Blakiston Fan is substantially lower than all other *S. semiluna* occurrences, we can infer that the Alberta population exhibits niche divergence relative to the rest of species and is unique in its ecological/environmental associations (Neal et al. 2018; Campbell et al. 2022). We also compared how the values of variables that were most important in predicting *S. semiluna* occurrences varied between the Alberta population and all other occurrences. Important variables were determined using MaxEnt mean permutational importance threshold of 5%. We extracted values of these variables for Blakiston Fan and all other *S. semiluna* occurrences and conducted PCA on these values to see if the Alberta population clustered within or outside other occurrences.

## Results

### Reference genome sequencing and assembly

We assembled the first reference genome for *S. semiluna* (Blakiston Fan, Alberta, Canada) using a combination of PacBio HiFi long-read sequencing and Omni-C proximity ligation. The final assembly consisted of 1,246,961,827 bp organized into 86 scaffolds with an N50 of 56.2 MB and an N90 39.2 MB, respectively (Fig. 1). Additional assembly statistics, an off-target and microbial contigs blob plot, *k*-mer profiles, Hi-Rise contiguity plots, and a Hi-C contact map showing the high contiguity of this genome can be found in Supporting Information (Tables S1 and S2, Fig. S1, S2, S3, S4, S5, and S6). Twenty-three putative chromosomes, including the Z (identified using BUSCO genes; 52.8MB), range in length from 29.0 to 77.6 MB and together contain 99.5% of the total sequence length. Of the 5,286 BUSCO genes from the lepidoptera_odb10 database, 95.9%, 1.2%, and 0.5% were found in the reference genome to be complete, fragmented, and missing, respectively. Overall, this reference genome is highly contiguous and very complete. RepeatMasker analyses identified a total of 718.1 MB of repetitive sequence within the reference genome, comprising 57.59% of its total length. Categorization of this repetitive sequence into retroelements, DNA transposons, rolling-circles, small RNA, satellites, and simple repeats is reported in Supporting Information (Table S3).

**Figure 1.**
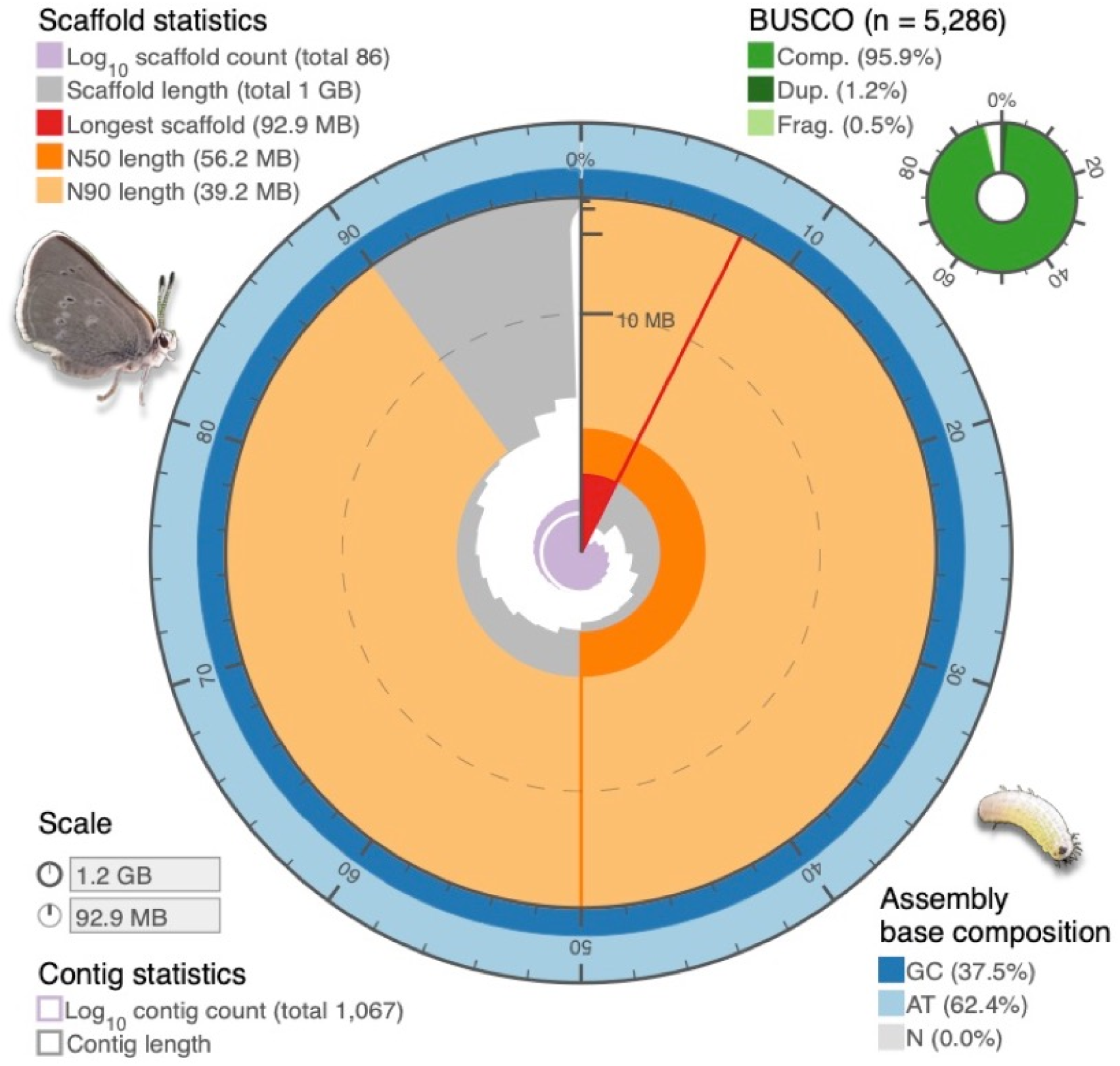
Snail plot showing quality metrics for the *Satyrium semiluna* genome assembly (Challis et al. 2020). The circle’s circumference represents the size of the assembly (∼1.2 GB). The radius represents the longest scaffold (with its length indicated by the red wedge relating to the circumference) measured on a log scale (with 10 MB tick on a vertical axis) and N50 and N90 metrics are displayed in different colors. N50 and N90 measure the length of the shortest scaffold in the group of the longest scaffolds that together contain >50% and >90% of the total genome length and are shown in dark and light orange arcs, respectively. Other scaffolds are ordered by size moving clockwise and drawn in gray. The central light purple spiral represents the cumulative scaffold count. The dark and light blue area around the outside of the circle represent the GC and AT content, respectively, at 0.1% intervals. Of the 5,286 BUSCO genes from the lepidoptera_odb10 database, 95.9%, 1.2%, and 0.5% were found to be complete, fragmented, and missing, respectively (Manni et al. 2021). Adult and larval individuals from the endangered population at Blakiston Fan, Alberta, Canada, are pictured (photographs by □RNG). This snail plot was generated using BlobToolKit (https://github.com/blobtoolkit/blobtoolkit)

### Sequence processing and genotyping

We completed whole-genome resequencing of 15 *S. semiluna* individuals on an Illumina NovaSeq platform, aiming for a mean coverage of ∼20x per sample. Across all individuals, a total of >1.4 billion paired-end, 150-bp reads were sequenced, passed Illumina filters, and were aligned to our *S. semiluna* reference genome. Sequence coverage of individuals ranged from 14.2x to 28.1x, with a mean of 19.1x (see Supporting Information Table S4 for reads per individual.) Joint genotyping resulted in a multisample VCF file consisting of 41,083,914 variants across the 15 individuals and a final post-filtering VCF file containing 23,889,641 SNPs. This post-filtering dataset was used in all subsequent analyses unless otherwise specified.

### Population structure

Pairwise *F*_ST_ values indicated substantial differentiation among the Alberta, British Columbia, and Montana populations: Alberta *vs*. British Columbia = 0.424; Alberta *vs*. Montana = 0.292; British Columbia *vs*. Montana = 0.322. There was no measurable divergence between individuals collected from the two British Columbia locations (∼17 km apart), indicated by a negative *F*_ST_ estimate (−0.004) that should be functionally interpreted as zero. Estimates of *d_xy_* aligned with those of *F*_ST_: Alberta *vs*. British Columbia = 0.009; Alberta *vs*. Montana = 0.005; British Columbia *vs*. Montana = 0.008.

Using a thinned set of 108,283 physically unlinked SNPs, PCA cleanly separated Alberta, British Columbia, and Montana individuals into discrete clusters (Fig. 2, panel a). PC1, which accounts for over a third of the total genomic variation, separated populations from west (British Columbia) and east (Alberta, Montana) of the Rocky Mountains/continental divide, PC2 (11% of the variation) separated Alberta and Montana, and individuals from British Columbia were spread along PC3 (6.5%) generally according to collection location. We further quantified population structure and degree of admixture using the model-based clustering program structure (Pritchard et al. 2000). Our first structure analysis of all individuals identified an optimal value of *K* = 2, with Alberta/Montana individuals forming one group and British Columbia individuals a second, with essentially no admixture. Subsequent runs including only Alberta and Montana individuals identified an optimal value of *K* = 2, separating the populations into two distinct clusters, again with virtually no admixture.

**Figure 2.**
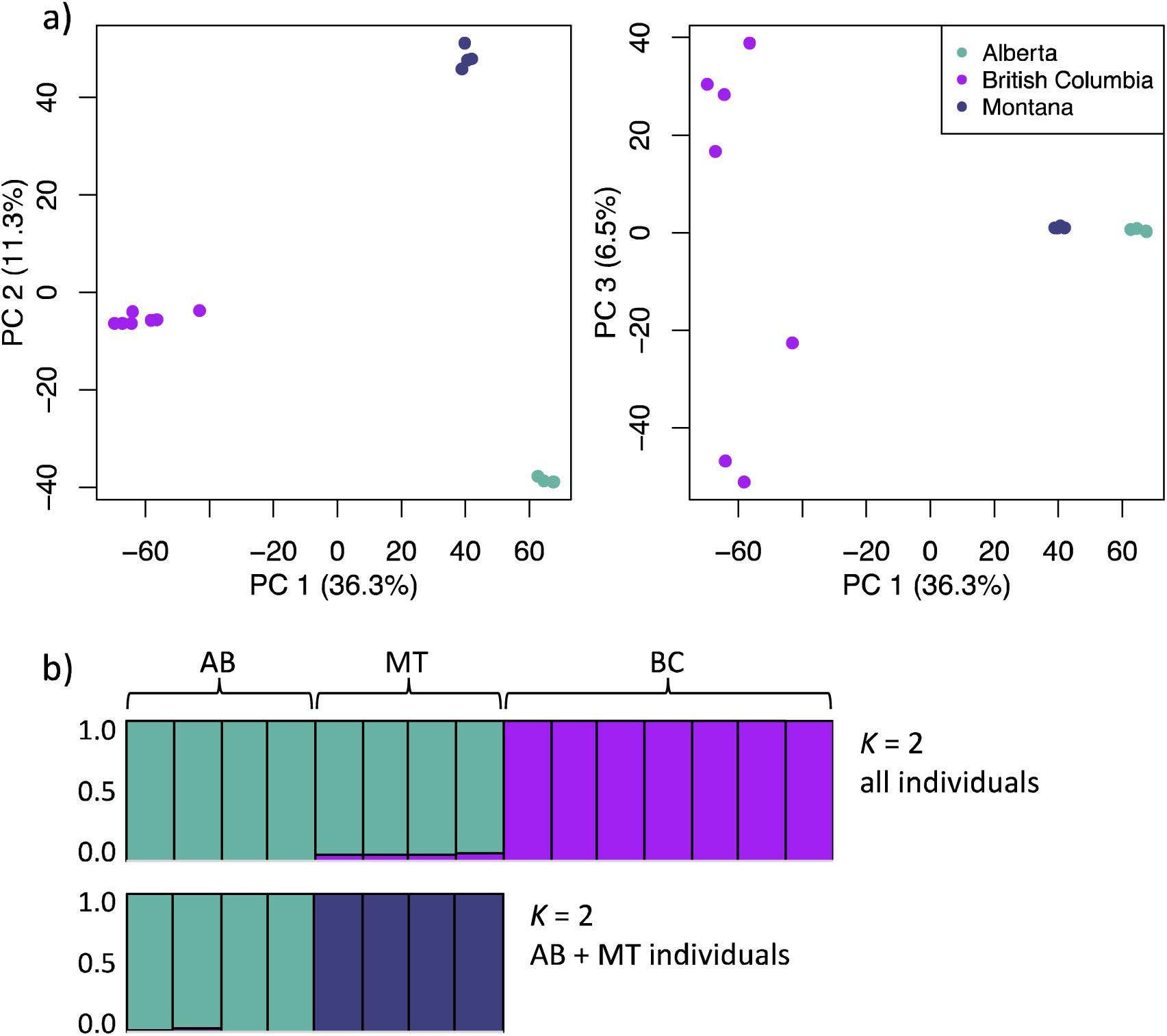
Population genetic structure of *Satyrium semiluna*, using a) principal component analysis (PCA), and (b) model-based clustering with the program structure. For PCA, every point represents a sequenced individual (*n* = 15), color coded according population. Our structure analyses addressing *K* = 1:4 found an optimal value of *K* = 2 when all individuals were analyzed together, splitting Alberta and Montana from British Columbia. Hierarchical runs of the Alberta and Montana individuals together identified an optimal value of *K* = 2 with virtually no admixture.

### Genetic diversity and runs of homozygosity

Heterozygosity for Alberta individuals (mean = 0.083) was roughly one third that of British Columbia (mean = 0.216) and half that of Montana individuals (mean = 0.154) (Fig. 3 panels a and b). Dividing the total number of heterozygous sites per individual by the total number of loci sequenced gives the following means per population: Alberta = 0.0016; British Columbia = 0.0040, and Montana = 0.0030. Heterozygosity within populations was generally consistent across individuals, as expected (Supporting Information, Fig. S7). Similarly, population π suggested that nucleotide diversity was far lower in the Alberta population (0.003) than both the British Columbia (0.008) and Montana populations (0.005).

**Figure 3.**
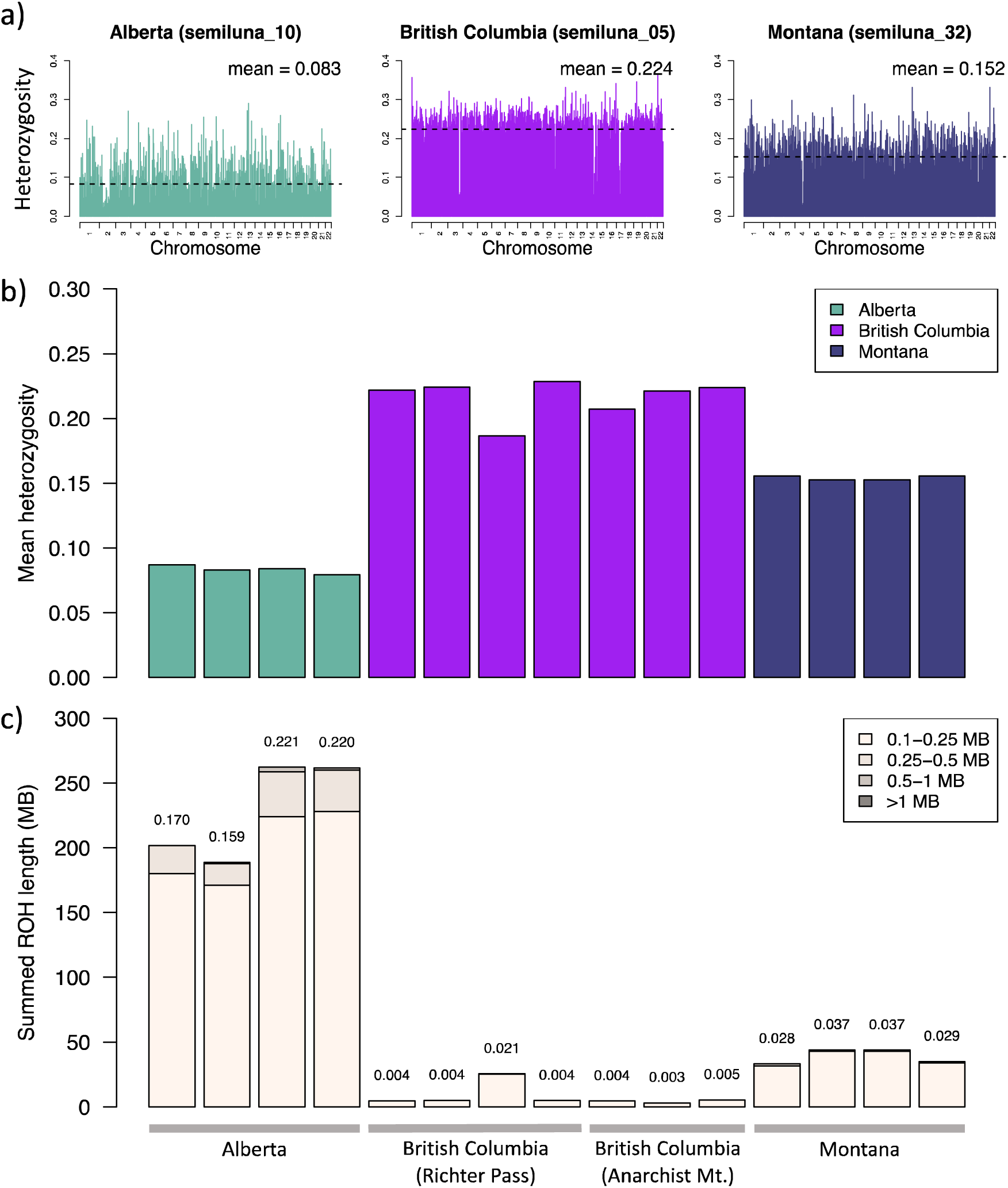
Analyses of zygosity in *Satyrium semiluna.* a) Heterozygosity for three example individuals, estimated as the average of heterozygous/homozygous loci within nonoverlapping 1 MB windows across the autosomal genome. The first individual (teal) is from an endangered population in Waterton Lakes National Park, Alberta, Canada, isolated from all other populations by >350 km. The second individual is from British Columbia, Canada (purple), and the third from Montana, USA (dark blue). b) Mean heterozygosity for all sequenced individuals (*n* = 15), estimated by averaging values across all 1 MB windows. Dividing the total number of heterozygous sites per individual by the total number of loci sequenced gives the following mean estimates per population: Alberta = 0.0016; British Columbia = 0.0040, and Montana = 0.0030. c) Cumulative length of runs of homozygosity (ROH) for each individual, categorized as short (0.1 – <0.25 MB), medium (0.25 – <0.5 MB), long (0.5 – <1 MB), or very long (>1 MB). An estimate of inbreeding, *F*_ROH_, estimated as the total length of all ROH calls >0.1 MB divided by the cumulative length of the autosomal genome, is reported above each individual’s bar. This statistic approximates the proportion of the genome for which both copies of region are identical by descent, a product of either recent inbreeding (long ROH) or historical inbreeding (short ROH).

ROH were observed to be 5 – 70 times more prevalent in Alberta individuals than British Columbia and Montana individuals (Fig. 3 panel b). This indicates that that inbreeding has been far more prevalent in the Alberta population (mean *F*_ROH_ = 0.192) than in British Columbia (mean *F*_ROH_ = 0.006) or Montana (mean *F*_ROH_ = 0.033) populations. In Alberta individuals, short ROH (0.1 – <0.25 MB) were most common, with some medium ROH (0.25 – <0.5 MB) and very few long ROH (0.5 – <1 MB) present. This is consistent with a long history of high inbreeding. However, no very long ROH (>1 MB), which are generally interpreted as evidence of recent inbreeding (i.e., within the past ∼10 generations), were present. In contrast, British Columbia and Montana individuals had far fewer ROH overall, and almost all were short, suggesting little or no historical or contemporary inbreeding within these populations. A second inbreeding coefficient provided similar relative rates of inbreeding, with the Alberta population far more inbred (mean *F*_HOM_ = 0.622) than British Columbia (mean *F*_HOM_ = 0.019) and Montana (mean *F*_HOM_ = 0.305).

### Demographic history and effective population size

PSMC showed that the three populations differed substantially in their demographic history, particularly in the more recent past (Fig. 4). From 2.5 mya to about 250 kya, all populations shared a similar demographic history of gradually increasing population sizes. At 250 kya, British Columbia continued a rapid, two-fold increase in *N*_e_ until 100 kya, when *N*_e_ began declining until 30 – 40 kya. Around 150 kya, the Alberta population, and to a lesser extent the Montana population, experienced an increase in *N*_e_, followed by a sharp decline at 100 kya until 30 – 40 kya. From 30 – 40 kya onwards, British Columbia and Montana both experienced increases in *N*_e_, while the Alberta population flatlined between 1,000 and 2,000 individuals from 40 – 5 kya. The sharp increase in *N*_e_ from ∼10 kya onwards suggested by multiple British Columbia individuals likely indicates that populations occurring west of the continental divide rapidly expanded and experienced broad scale connectivity following the end of the Younger Dryas (Clement and Peterson 2008). PSMC curves did not systematically differ between individuals collected from the two British Columbia locations.

**Figure 4.**
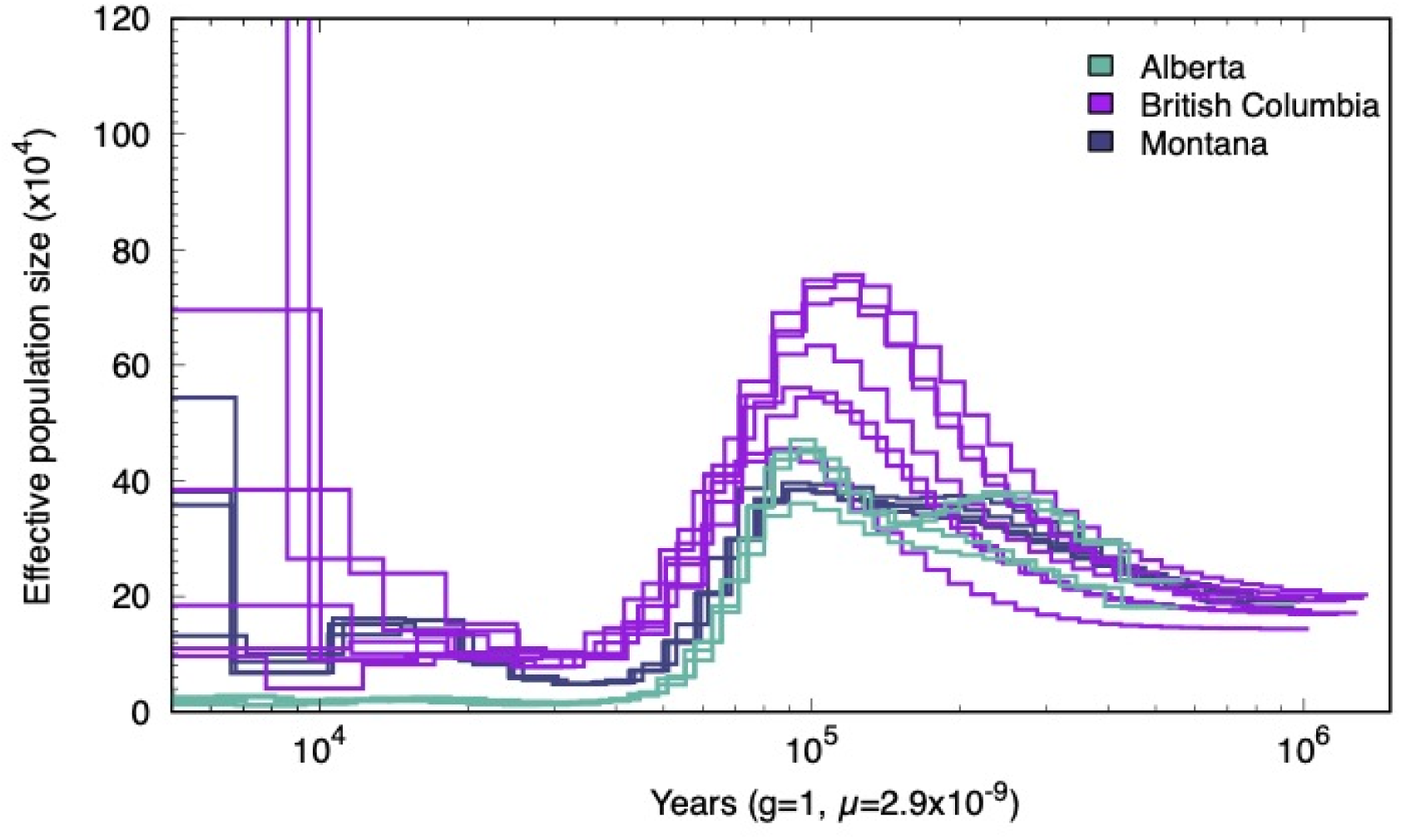
Pairwise Sequentially Markovian Coalescent (PSMC) analyses from 2.5 mya until 5 kya of three *Satyrium semiluna* populations using whole-genome data. The estimated effective population size (*N*_e_) is shown on the y-axis and coalescent time in years before present on the x- axis. Four individuals (teal lines) are from an endangered population in Waterton Lakes National Park, Alberta, Canada, isolated from all other populations by >350 km. Seven individuals are from two locations in British Columbia, Canada (purple lines), and four individuals from a single location in Montana, USA (dark blue lines). Most interestingly, from 30 – 40 kya onwards, British Columbia and Montana both experienced increases in *N*_e_, while the Alberta population flatlined between 1,000 and 2,000 individuals from 40 – 5 kya.

To estimate contemporary *N*_e_, we used currentNe (Santiago et al. 2023), which relies on linkage disequilibrium among SNPs for multiple individuals sampled from a single population. Estimates of contemporary *N*_e_ using LD between pairs of SNPs located on different putative chromosomes varied widely among populations. In all cases, currentNe converged and provided estimates of *N*_e_ with both 50% and 90% confidence intervals:

Alberta *N*_e_ = 487.11; 50% CI = 154.57, 1,535.05; 90% CI = 29.65, 8,001.68

British Columbia *N*_e_ = 755.20; 50% CI = 270.17, 2,111.01; 90% CI = 61.53, 9,264.72

Montana *N*_e_ = 14920.04; 50% CI = 2503.76, 88,909.16; 90% CI = 191.97, 11,160,397.45

### Niche analyses and prediction of suitable habitat

Our ecological niche models predicted *S. semiluna* occurrences with a high accuracy, reflected in a mean AUC score of 0.94. Suitable habitat was relatively sparse within the central and northern range of *S. semiluna* (Fig. 5, panel a). The relative contribution of each environmental variable, estimated as both mean permutational importance and percent contribution (we favor permutation importance, see Searcy and Shaffer 2016), are reported in Supporting Information Table S5. Overall, variation in precipitation variables were most important in predicting *S. semiluna* occurrences, with temperature and landscape/geographical variables generally less important. Mean summer precipitation (June to August) was the single most important variable, with mean permutation importance and percent contribution more than double that of mean autumn precipitation (September to November), the next most important variable.

**Figure 5.**
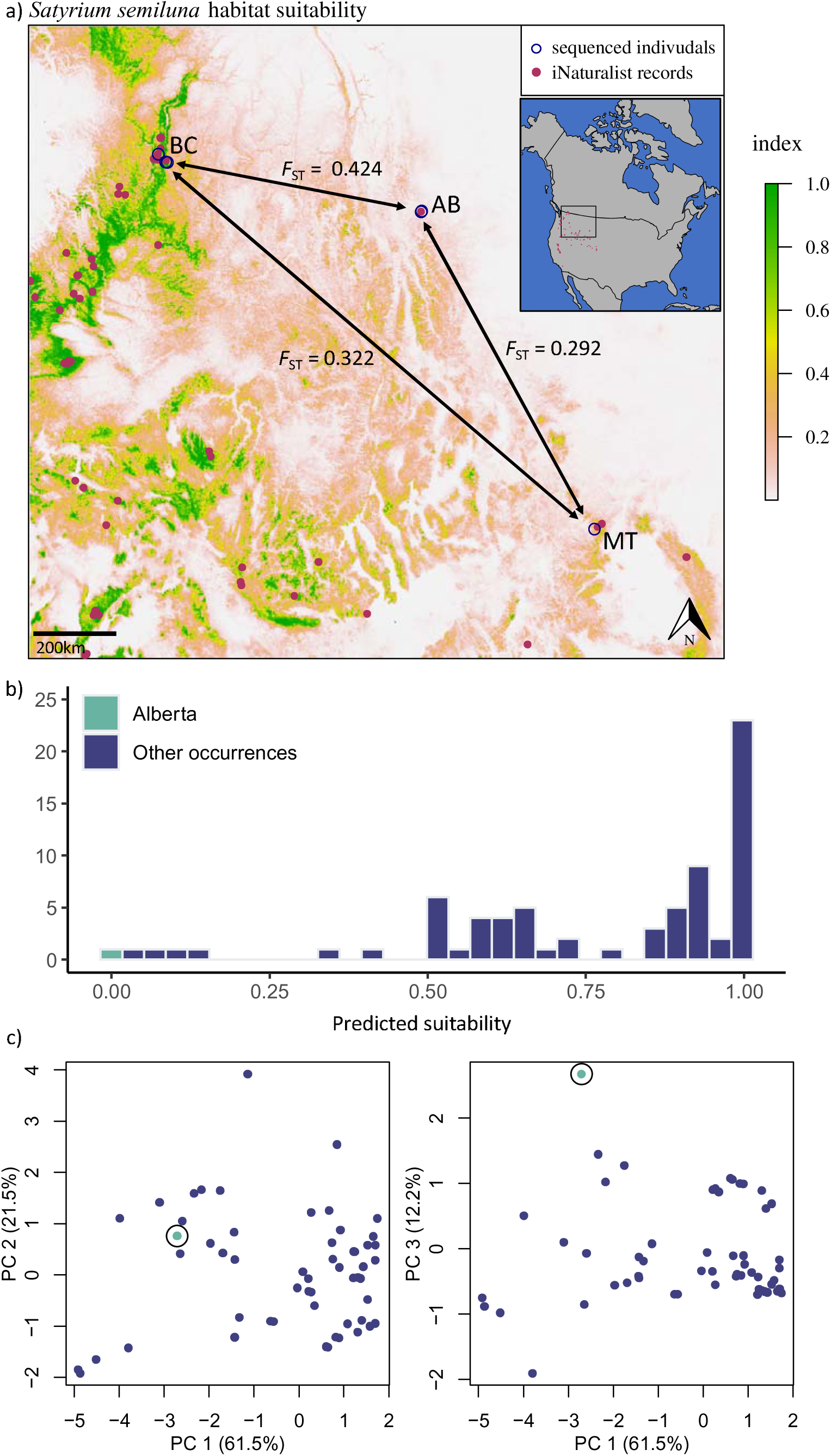
a) Predicted habitat suitability across the central and northern extent of the range of *Satyrium semiluna*, with higher index scores corresponding to higher suitability. Vetted iNaturalist occurrences and the collection locations of sequenced individuals were used to parameterize this model. Predictors included both environmental/ecological and geographical variables. b) Predicted habitat suitability values for all iNaturalist observations and the location of the Alberta population at Blakiston Fan. These predicted values were generated using a model identical to the one above, except all Alberta locations were removed for parameterization. Using this model, Blakiston Fan had a predicted suitability value of 0.003, indicating the ecological and environmental conditions at this site are very atypical for the species. c) Principal component analysis (PCA) on the extracted values of all predictor variables with a mean permutational importance above 5% in the first model. Ecological and environmental values of the Alberta population did not fall within the range of other occurrences on PC3, for which mean summer precipitation had the highest loading.

Excluding all occurrences of the Alberta population resulted in greater predictive power (AUC = 0.97) and a predicted habitat suitability surface that was very similar to the first (Pearson *r* = 0.94). Using this model, predicted suitability for all non-Alberta occurrences remained high (mean = 0.77, s.d. = 0.27) while that of the Alberta population was essentially zero (Blakiston Fan = 0.003) (Fig. 5, panel b). Four other *S. semiluna* occurrences had very low predicted suitability values (< 0.15), as would be expected for any semi-vagile species that may occasionally wander outside of suitable habitat. Values of all variables with a mean permutational importance above 5% were very dissimilar between these low suitability occurrences and the Alberta population, indicating that they are not ecological/environmental matches. The predicted habitat suitability value of Blakiston Fan was 0.003—essentially zero. This suggests that the Alberta population is associated with ecological and environmental conditions that are atypical of the species, equating to niche divergence. PCA on the extracted values of all variables with a mean permutational importance above 5% indicates that ecological and environmental associations of the Alberta population did not fall within the range of other occurrences, again consistent with strong niche divergence (Fig. 5, panel c). This divergence was apparent on PC 3, for which mean summer precipitation had by far the highest loading.

## Discussion

Genetic rescue is a viable conservation practice that can ameliorate genetic erosion, inbreeding depression, and loss of adaptive potential in small and isolated populations (Ingvarsson 2003; Pimm et al. 2006; Trinkel et al. 2008; Finger et al. 2011; Mattila et al. 2012; Mussmann et al. 2017; Frankham et al. 2017; Clarke 2024). In most outbreeding species, net-positive effects of enhanced population connectivity and gene flow have become a null expectation (Frankham et al. 2017). We generally support this view, particularly when considering populations recently fragmented through human activities. Still, conservation practitioners have been reluctant to implement genetic rescue, both because of inherent risks (and associated responsibility) involved in active interventions, and because genetic/genomic factors are often simply ignored in conservation planning (Shafer et al. 2015; Frankham et al. 2019; Liddell et al. 2021). In this study, we highlight how whole-genome data can be quickly generated for an endangered species with no previous genomic resources, and used to evaluate conservation strategies, including genetic rescue.

Key data necessary for this assessment included a well-assembled reference genome and whole-genome sequence data for both recipient and potential donor populations. We produced the first high-quality reference genome for *S. semiluna* and the subfamily Theclinae. At 1.25 GB, this genome is almost double the size of the next largest scaffold- or chromosome-level genome available for the family Lycaenidae, and one of the largest for any butterfly (further discussion on genome size in Supporting Information). Our analyses of millions of SNPs across the genomes of 15 individuals confirm that the Alberta population is very divergent from the geographically nearest populations in British Columbia (*F*_ST_ = 0.40) and Montana (*F*_ST_ = 0.28). Genetic diversity, defined here as the mean of individual heterozygosity, was on average 2.5 times higher in British Columbia individuals and 1.9 times higher in Montana individuals than Alberta individuals. Runs of homozygosity (ROH) were 30.4 and 5.9 times more prevalent in Alberta individuals than British Columbia and Montana individuals, and inferences of inbreeding based on *F*_ROH_ suggest that that more than 20% of Alberta individuals’ genomes were devoid of SNP diversity and identical by descent. Overall, the Alberta population shows signs of genomic flatlining (Robinson et al. 2016; Mooney et al. 2023) and a high level of inbreeding, similar to observations in Channel Island foxes (Robinson et al. 2016; 2018), Ethiopian wolves (Mooney et al. 2023), Isle Royale gray wolves (Kardos et al. 2018b; Robison et al. 2019), Isle Royale moose (Kyriazis et al. 2023), and Muskox (Pečnerová et al.2023). (See Supporting Information Fig. S8 for ROH estimates with repetitive sequences masked.) However, unlike many of these large mammal populations, the absence of long ROH (e.g., >1Mb) in the Alberta *S. semiluna* population suggests that reduced genetic diversity is primarily a product of a long history of inbreeding, rather than a recent, sustained bottleneck (Stoffel et al. 2021). A somewhat similar observation was made for Ethiopian wolves, consistent with a long history of small population sizes (Mooney et al. 2023). This information, combined with the absence of admixture among populations (Fig. 2 panel b), suggests that the Alberta population has been both small and isolated for a very long time.

Whether small and isolated populations will persist without intervention, or if genetic processes will lead to their inexorable extinction, is largely thought to hinge on contemporary population size (Franklin 1980; Soulé 1980; Lande and Barrowclough 1987; Frankham et al. 2014; 2017). We estimated the contemporary *N*_e_ of the Alberta population to be approximately 487 individuals based on linkage disequilibrium among SNPs in four individuals. Repeat transect surveys completed in 2022 and 2023 resulted in estimates of 8.1 and 7.9 adult individuals per hectare, respectively. Assuming all ∼300 hectares of Blakiston Fan are uniform in habitat quality yields a census estimate of 2430 and 2370 individuals, respectively. While an assumption of habitat uniformity is likely invalid, these estimates fall within the low end of the qualitative historical population estimate of 2,000 – 10,000 individuals (COSEWIC 2006). Our estimate of *N*_e_ is lower than survey population estimates, which is expected because not all individuals contribute equally to the breeding pool (Frankham 1995). Between the two measures, *N*_e_ is generally regarded as the more important statistic for conservation management. For example, according to the widely debated 50/500 rules, a minimum *N*_e_ of 50 individuals is necessary to avoid inbreeding and 500 individuals to avoid potentially deleterious levels of genetic drift (Franklin 1980; Soulé 1980; Lande and Barrowclough 1987). Based on these thresholds, the Alberta population could theoretically maintain its genetic diversity in isolation. However, modern theory and evidence suggest that a minimum *N*_e_ of 100 individuals is required to limit inbreeding depression to 10% over 5 generations and 1000 individuals for retaining evolutionary potential (Frankham et al. 2014). We might therefore doubt that the Alberta population is of sufficient size for long-term persistence. However, field observations confirm that the Alberta population not only continues to persist, but is also capable of rapid recovery from population crashes, as was observed within the last decade.

It is generally assumed that over time, genetic diversity in small and isolated populations will decline, and the benefit from genetic rescue will increase (Frankham et al. 2017; Ralls et al. 2018). However, as the aforementioned authors point out, this conclusion is drawn from plant and animal populations fragmented within the last ∼500 years (Frankham et al. 2017; Ralls et al. 2018). When considering populations recently fragmented from anthropogenic activities, we might expect successful purging to be exceedingly rare, because any population that becomes small and isolated would need to survive the substantial mortality associated with recessive variants drifting to high frequency and being culled from the population in selection against homozygous individuals. Therefore, for these populations, facilitating gene flow via genetic rescue is almost always a better conservation strategy than holding out for genetic purging (Ralls et al. 2020). However, the process of genetic purging may still be an important consideration in conservation. Many small extant populations with a very long history of isolation (i.e., fragmented well before widespread anthropogenic activities) likely represent those rare survivors that successfully purged a large portion of their deleterious recessive alleles, aiding in their persistence and rendering present-day inbreeding less problematic (Robinson et al. 2016; 2018; 2022; Pečnerová et al. 2024).

Our PSMC analyses suggest that the Alberta *S. semiluna* population has been both small (<2000 individuals) and completely isolated from 5k – 40k years before present. Due to the relatively small number of individuals sequenced, and lack of phased genomic data and a recombination map, inferring more recent changes in historical *N*_e_ is not possible (Beichman et al. 2018; Barroso et al. 2019; Nadachowska-Brzyska et al. 2022; Gargiulo et al. 2024). Increased sampling of populations may result in more robust estimates of contemporary *N_e_* and facilitate additional analyses to compare to those employed here. For example, the high *N_e_* estimates for the Montana population were surprising, and deserve further scrutiny. This remains an important knowledge gap that future research should address. Still, we can draw several meaningful inferences from our historical *N*_e_ analyses, which encompass the last glacial maximum (20 – 26 kya). During this period, the Rocky Mountain Foothills of Alberta, adjacent to Blakiston Fan, were likely glaciated by a continental (Laurentide) ice sheet (Jackson and Little 2004). It is unclear the extent to which the area now known as Blakiston Fan was glaciated. However, southwestern Alberta and the surrounding area was thought to have been rich in small and isolated glacial refugia (Dyke et al. 2003; Shafer et al. 2010). Based on both patterns of genetic differentiation and PSMC analyses, we infer that at either Blakiston Fan or another locality east of the continental divide, a small population of *S. semiluna*, with descendants now occupying Blakiston Fan, persisted in isolation through the last glacial maximum. We observed no evidence that this isolated population had any gene flow with populations to the south, beyond the Laurentide ice sheet, indicated by both its consistently small *N*_e_ through this period and lack of admixture with other populations. Based on this best-available information, we infer that historical gene flow has not played a role in long-term persistence of the Alberta population.

Long histories of isolation have important consequences not only for patterns of genetic diversity and differentiation, but also ecological diversity and differentiation. We hypothesized that differences in local environmental conditions coupled with complete isolation led to niche divergence of the Alberta population. Our analyses of niche divergence using ecological niche modelling confirm that the Alberta population exhibits environmental associations that are very atypical of the species. Mean summer precipitation, the most important variable for predicting occurrences of *S. semiluna*, was not highly correlated with other predictor variables included within our niche models (|*r*| < 0.70). PCA indicated that this variable also contributed the most to differentiating the Alberta population from all others. Based on ClimateNA, Blakiston Fan receives a total average rainfall of ∼200 mm over the summer months, while other *S. semiluna* occurrences included in our analyses range from 32 to 154 mm (mean = 71 mm). Four other occurrences observed to exhibit low suitability (Fig. 3 panel b) range in summer precipitation from 55-120 mm, indicating they are not similar to Blakiston Fan. Whether these observations represent actual populations or vagrant individuals is unclear and requires future investigation. Regardless, based on the observed associations, we infer that the Alberta population is uniquely adapted to substantially wetter summers relative to all other populations in the central and northern extent of the species’ range. Based on two of the four criteria provided by Frankham et al. (2011)—populations have exchanged no genes in the last 500 years and inhabit different environments—we predict that outbreeding depression would be a likely outcome if the Alberta population was crossed with others. We hope that this prediction can be tested experimentally with *ex situ* population crosses to quantitatively evaluate population compatibility and the fitness of hybrid offspring.

In 2021, 2022, and 2023, surveyors searched for *S. semiluna* larvae on both *L. argenteus* and *L. sericeus* at Blakiston Fan with approximately equal effort, searching about 10,000 individual plants per species per year. A total of 380 larvae were observed over those three years, all on *L. argenteus* and none on *L. sericeus* (unpublished data). While we have not completed ovipositional choice experiments, it is reasonable to infer that females would at least occasionally oviposit on *L. sericeus* in nature if it was at all preferred, given its availability at Blakiston Fan. In contrast, in British Columbia, *L. sericeus* is the only lupine species available in *S. semiluna* habitat and it is readily used by populations there (unpublished data). In Montana, both *L. argenteus* and *L. sericeus* have been reported in *S. semiluna* habitat on iNaturalist, suggesting that both species are available. However, host associations of these populations are currently unknown and require investigation. Representing yet another possible axis of niche divergence, we (J.R.N.G.) recently noted that larvae of the Alberta population share a mutualistic relationship with a single species of ant, *Lasius ponderosae*. In British Columbia, we observed that *L. ponderosae* is not present in *S. semiluna* habitat, and that *S. semiluna* larvae instead associate with *Formica* sp. and *Camponotus* sp.. Populations in California have been observed to associate with these genera as well (Runquist 2012). The importance of these relationships (obligate *vs.* facultative), as well as the extent and specificity of myrmecophilia across the range of *S. semiluna*, is not known, but may be important for any considerations of possible genetic rescue.

Our recommendation at this time is that genetic rescue is an inappropriate strategy for the conservation of the Alberta *S. semiluna* population. However, this recommendation is contingent on the continued absence of substantial inbreeding depression. In another butterfly, *M. cinxia*, inbreeding depression in a small and isolated island population was observed in the form of reduced egg viability (Mattila et al. 2012). This was alleviated when inbred individuals were hybridized with those of other, genetically diverse populations, demonstrating that genetic rescue should be an important part of recovery strategies for butterflies. However, it is important to note that this island *M. cinxia* population and Alberta *S. semiluna* population differ in their duration of isolation by many orders of magnitude (∼75 *vs.* up to 40k years). Reduced fitness attributed to inbreeding depression within the *M. cinxia* island population is evidence that genetic purging had not occurred, at least for the traits investigated, and that the population was much more likely to benefit rather than suffer from genetic rescue interventions. Inbreeding depression has never been explicitly investigated in *S. semiluna*, but we have several lines of evidence to suggest that, at present, it does not negatively impact population persistence. Even when the estimated population density plummeted to 1.5 individuals/ha in 2017 and 0.5 individuals/hectare in 2018, the population rapidly recovered to pre-crash levels in just a few years, peaking at 30.6 individuals/ha in 2021. If the Alberta population’s pre-crash inbreeding load was substantial, this bottleneck would have resulted in a high genetic load and reduced population recovery (Roelke et al. 1993; Westemeier et al. 1998). Such an inability to recover from bottlenecks is a defining property of populations that suffer from inbreeding depression (Frankham et al. 2017), and there was no indication of it based on extensive field surveys.

Genetic rescue is a proven means for ameliorating inbreeding depression, bolstering genetic diversity, and restoring individual fitness and population viability in the majority of circumstances (Frankham et al. 2011; 2017; 2020), and our conclusions from this study should not be interpreted as a recommendation that it be downgraded as a management strategy. Deliberate augmentation of gene flow and mixing of lineages are inevitabilities if we wish to preserve populations fragmented through recent anthropogenic activities. Our primary emphasis here is on biological outcomes—if genetic rescue will enhance survivability, we endorse it. However, if interventions are not needed and may be harmful, as in the case under consideration here, we hope managers will proceed with caution. In addition, we note that a desire to maintain ecological or evolutionary purity or identity also comes into play. To guide its decision-making process, Parks Canada implements a concept of “ecological integrity”, defined as “a condition that is determined to be characteristic of its natural region…” (Canada National Parks Act 2000). If the genetic composition of a particular population is assessed to be characteristic of a natural region, then genetic rescue may be viewed as a threat to ecological integrity (MacDonald et al. *In Review*). However, aiming above all else to preserve the purity/identity of certain evolutionary lineages risks not only their extinction, but also that of more inclusive evolutionary significant units, including entire species. We argue that evolutionary or ecological divergences among populations or lineages should only affect decisions to attempt genetic rescue insofar as they influence population fitness and viability. Parmesan et al. (2023) convey that evolutionary biologists are generally partial to hybridizations among divergent populations, recognizing a large set of benefits. In contrast, conservation practitioners often oppose intentional admixture because it jeopardizes the purity/identity of existing groups. Even within our *S. semiluna* conservation working groups, we observe this divergence in perspectives. Both are valid, and the tension between these two schools of thought is as much about conservation aesthetics as it is conservation biology. We hope that this case study on *S. semiluna* contributes to evidence-based evaluations of genetic rescue, considering multiple perspectives, and prioritizing population and species persistence.

The genomic and ecological analyses presented here led to the designation of the Alberta *S. semiluna* population as a distinct designatable unit under Canada’s Species at Risk Act, and support management without genetic intervention. However, flatlined genetic diversity likely translates to low adaptive potential, and climate change is a harsh reality. If the Alberta *S. semiluna* population is observed to be maladapted to changing environmental conditions, intentional admixture with other *S. semiluna* populations may introduce beneficial genetic variation and bolster adaptive potential (Ralls et al. 2020). As with all endangered species, the need for genetic rescue must be continuously evaluated, both as new information becomes available and as circumstances change.

## Author contributions

All authors collectively conceived of the study design. Fieldwork was completed by JRNG. Lab work was completed by ZGM and FAHS. Reference genome curation and individual genotyping were completed by ZGM and JRD. Analyses were completed by ZGM with input from all authors. Writing was completed by ZGM with assistance from all authors.

## Acknowledgments

We wish to acknowledge Lacey Hébert and Llewellyn Haines for fieldwork support, Steve Kohler for providing Montana specimens, Natasha Lloyd for logistical and permit support, Janet Sperling for assistance with lab work, including DNA barcoding of specimens to confirm species-level identifications, Eric Runquist for consultation, members of the UCLA Shaffer lab for analysis comments and critiques, the Waterton Lakes National Park Ecological Restoration Team for leading habitat restoration programs and assisting population monitoring, and the Half-moon Hairstreak Conservation committee for discussions on impacts of this research. Funding was provided by the Calgary Zoo Foundation, Parks Canada (GC-1341), Shell Canada, a Natural Sciences and Engineering Research Council (NSERC) Discovery Grant awarded to FAHS (RGPIN-2018–04920), and a La Kretz Center for California Conservation Science Postdoctoral Fellowship and NSERC Postdoctoral Fellowship (PDF - 578319 - 2023) awarded to ZGM.

## Data Availability Statement

All genomic short-read data produced in this study will be made available on NCBI (SRA: SUB14903224). Our reference genome for Satyrium semiluna is available on NCBI (BioProject PRJNA907836). All spatial data layers used in this study will be made available on Dryad (doi:XX).

