## Supporting Information for "Whole-genome evaluation of genetic rescue: the case of a curiously isolated and endangered butterfly"

Reference genome sequencing and assembly

Four of the eight Alberta individuals were used to generate a chromosome-level reference genome for *Satyrium semiluna*. These individuals were euthanized by flash freezing and stored at -80°C. Reference genome library preparation (PacBio HiFi and Dovetail Omni-C) and sequencing was conducted by Dovetail Genomics. High molecular weight DNA was extracted from flash-frozen thoracic tissue from two male individuals using a Nanobind Tissue Big DNA kit. These samples were pooled and used to construct a HiFi SMRTbell library using the SMRTbell Express Template Prep Kit v2.0. A total of 2,417,793 PacBio CCS reads (29.1 Gb) were used as an input to Hifiasm v0.15.4-r347 (Cheng et al. 2021), run with default parameters. Blast results of the Hifiasm output assembly against the nt database were used as input for blobtools v1.1.1 (Laetsch & Blaxter 2017) and scaffolds identified as possible contamination were removed from the assembly. Finally, purge_dups v1.2.5 (Guan et al. 2020) was used to remove haplotigs and contig overlaps.

For the Dovetail Omni-C library, chromatin was fixed in place with formaldehyde in the nucleus and then extracted. Fixed chromatin was digested with DNAse I, chromatin ends were repaired and ligated to a biotinylated bridge adapter followed by proximity ligation of adapter containing ends. After proximity ligation, crosslinks were reversed and the DNA purified. Purified DNA was treated to remove biotin that was not internal to ligated fragments. Sequencing libraries were generated using NEBNext Ultra enzymes and Illumina-compatible adapters. Biotin-containing fragments were isolated using streptavidin beads before PCR enrichment of each library. The library was sequenced on an Illumina HiSeqX platform to produce approximately 30X sequence coverage.

The BlobToolKit pipeline was used to identify and remove off-target microbial contigs using nucleotide blast (Camacho et al. 2009) and protein DIAMOND BLAST (Buchfink et al. 2021, Buchfink et al. 2015) (Figure S1). General genome characteristics were assessed from *k*-mer distributions of debarcoded raw reads which were estimated with Jellyfish v2.2.3 (Marçais and Kingsford 2011) and GenomeScope v1.0 (Vurture et al. 2017) (k = 21, k-mer coverage cutoff = 10,000) (Figure S2). To ensure the assembly was largely free of diploid content, KAT v2.4.1 (Mapleson et al. 2016) was used in comp mode (*k*=27 and otherwise default settings) to compare *k*-mer spectra of debarcoded raw reads to those of the assembly (Figure S3).

The Hifiasm output assembly and Omni-C library reads (MQ>50) were used as input data for HiRise, a software pipeline designed specifically for using proximity ligation data to scaffold genome assemblies (Putnam et al. 2016). Dovetail OmniC library sequences were aligned to the draft input assembly using bwa (Li and Durbin 2009). The separations of Dovetail OmniC read pairs mapped within draft scaffolds were analyzed by HiRise to produce a likelihood model for genomic distance between read pairs, and the model was used to identify and break putative misjoins, to score prospective joins, and make joins above a threshold (Figure S4 and S5). Heat map of final scaffolded assembly is provided in Figure S6.

Identification of the Z chromosome

Many population, conservation, and landscape genomic analyses depend on autosome-only data due to differences in *N_e_* and unique evolution of the sex chromosomes (Charlesworth et al. 1987). No other chromosome-level reference genomes are currently available for butterflies in the subfamily Theclinae, so we used synteny analysis across the most closely related available subfamilies to identify the Z chromosome (Polyommatinae, estimated divergence times from Theclinae: ~56-100 MYA (Pohl et al. 2009)). Typical synteny approaches relying on sequence-to-sequence mapping, such as MUMmer4 (Marçais et al. 2018) and minimap2 (Li 2018), failed to reliably identify homologous sequence due to high levels of sequence divergence. We therefore ran BUSCO v5.2.2 on chromosome-level genome assemblies for four other butterfly species (Polyommatinae: *Polyommatus icarus*, NCBI accession GCA_937595015.1; *Glaucopsyche alexis*; NCBI accession GCA_905404095.1; *Aricia agestis*, NCBI accession GCA_944452695.1; Nymphalidae: *Melitaea cinxia*, NCBI accession GCA_905220565.1), took the union of BUSCO genes that occurred on their Z chromosomes, removed fragmented and duplicated genes, and filtered the *S. semiluna* BUSCO output for these genes only. All 81 were located on a single scaffold (ScvBUXZ_10.HRSCAF233), which we inferred to be the Z chromosome.

Discussion on genome size

At 1.25 GB, this genome is almost double the size of the next largest scaffold- or chromosome-level assembly available on NCBI for Lycaenidae (728.8 MB, *Calycopis cecrops*, NCBI accession GCA_001625245.1). Currently (June 17, 2024), 23 scaffold- or chromosome-level assemblies are available for 16 species of lycaenid butterflies, with an average of 501.5 MB. No chromosome-level assemblies are available for the Theclinae, making this the first for the subfamily. Within Lycaenidae, Theclinae is reported to have the largest average genome size, with estimates ranging from 225–798 MB (Liu et al. 2020). The size of our *S. semiluna* assembly is well outside this range. The cause of this deviation in genome size is not known, but may be attributed to a high frequency of repetitive sequence (Petrov 2001; Liu et al. 2020). Variation in transposable elements, in particular, have been shown to play a major role in genome size variation within Lepidoptera (Talla et al. 2017). RepeatMasker analyses identified a total of 718.1 MB of repetitive sequence, or 57.59 % of the total genome length , of which, >95% were classified as transposable elements (Table SX). Baselines for the repetitive content of butterfly genomes are yet to be established, so we cannot be sure if this entirely accounts for the large genome size of *S. semiluna*. Lepidoptera also exhibit substantial variation is chromosome number, ranging from five to 226, the largest number of chromosomes observed in any non-polyploid eukaryotic organism (*Polyommatus atlanticus*) (Brown et al. 2004; Kandul et al. 2007; Saura et al. 2013; Lukhtanov 2014; 2015). However, we identified 23 putative chromosomes within our homogametic assembly (Fig. S6), suggesting that an increase in chromosome number is not a cause of the large genome size.

Pairwise Sequentially Markovian Coalescent (PSMC) analyses

Historical fluctuations in Ne were estimated from 2.5 Ma until 5 ka using Pairwise Sequentially Markovian Coalescent (PSMC) analyses (Li & Durbin, 2011). The primary input for PSCM is a consensus genome sequence (fastq) for each individual. We generated fastq files using individual-level BAM files (produced in genotype calling) and the mpileup command (-C 50, -Q 30, and -q 30) in samtools (Danecek et al. 2021) piped into the vcf2fq command from vcfutils.pl, with our S. semiluna genome assembly used as the reference. We filtered sites with inferred consensus quality <20 and a read depth less than 8× or greater than two times the each individual samples’ mean coverage, calculated from BAM files using samtools “depth”. When running PSMC, we set the maximum number of iterations (N) to 25, maximum coalescent time (t) to 5, initial theta/rho ratio (r) to 5, and parameter pattern (p) to “4 + 30*2 + 4 + 6 + 10”. We set the generation time to 1 year, as *S. semiluna* is known to be univoltine, and the mutation rate to 2.9 × 10-9 mutations per year, based on the best estimate for *Heliconius melpomene* (Keightley et al. 2015).

Niche analyses and prediction of suitable habitat

To parameterize MaxEnt models, we compiled 17 environmental variables of interest using ClimateNA v7.30 software (Wang et al. 2016; Mahony et al. 2022; downloaded via AdaptWest Project. 2022; see Supplementary Materials for details). These included mean annual temperature and precipitation, mean temperature and precipitation of the summer, fall, winter, and spring, the difference between mean temperatures of the warmest and coldest months (“continentality”), degree-days below 0°C (chilling degree days), degree-days above 5°C (growing degree days), mean annual precipitation as snow, mean annual relative humidity, and length of the frost-free period. MaxEnt models were parameterized using 10,000 background points, with all feature classes made available and the regularization parameter set to 1.0 (Phillips 2005). To evaluate predicative power, we withheld 20% of occurrences for cross-validation and receiver operating characteristic (ROC) analysis using five different models (Phillips et al. 2006). We then ran a full model with all occurrences to predict habitat suitability across the study landscape (logistic output) using the “predict” function (raster package). All 1 km grid cells received a habitat suitability score ranging from 0 – 1, with higher values indicating higher suitability.

Table S1. Statistics for both Hifiasm and HiRise assemblies.

|  | Hifiasm Assembly | HiRise Assembly |
| --- | --- | --- |
| Largest scaffold | 10,916,870 | 92,921,669 |
| Number of scaffolds | 960 | 86 |
| Number of scaffolds > 1kbp | 960 | 86 |
| Number of gaps | 43 | 932 |
| Number of N's per 100 kbp | 0.08 | 7.21 |

Table S2. Contiguity metrics of our final *Satyrium semiluna* reference genome.

| Assembly | Total Length (bp) | N50 | L50 | N90 | L90 |
| --- | --- | --- | --- | --- | --- |
| Input Assembly | 1,246,872,927 | 2,459,303 | 146 | 739,926 | 500 |
| Dovetail HiRise Assembly | 1,246,961,827 | 56,152,807 | 10 | 39,200,393 | 20 |

Table S3. Results of RepeatMasker analysis (v4.1.4; Smit et al. 2015), quantifyingrepetitive content, including transposable elements, endogenous retroviruses, and repeat motifs. A library of repetitive elements for Lepidoptera was compiled using Dfam v3.6 (Hubley et al. 2016). HMMER (nhmmscan v3.3.2; hmmer.org) was used to search the reference sequence against this library using default settings, opting to categorize simple repeats. Most repeats fragmented by insertions or deletions have been counted as one element. RepeatMasker analyses identified a total of 718.1 MB of repetitive sequence, or 57.59% of the total genome length.

| Classification | | Number of elements* | Length occupied (bp) | Percentage of sequence |
| --- | --- | --- | --- | --- |
| Retroelements: | | 1759055 | 383014481 | 30.72 |
|  | SINEs: | 577464 | 68773088 | 5.52 |
|  | Penelope | 43529 | 10999675 | 0.88 |
|  | LINEs: | 1068657 | 233785448 | 18.75 |
|  | CRE/SLACS | 19695 | 1907633 | 0.15 |
|  | L2/CR1/Rex | 209831 | 58358229 | 4.68 |
|  | R1/LOA/Jockey | 116607 | 47867975 | 3.84 |
|  | R2/R4/NeSL | 67307 | 8357947 | 0.67 |
|  | RTE/Bov-B | 218112 | 37384641 | 3 |
|  | L1/CIN4 | 679 | 31633 | 0 |
|  | LTR elements: | 112934 | 80455945 | 6.45 |
|  | BEL/Pao | 28517 | 35187592 | 2.82 |
|  | Ty1/Copia | 21088 | 3548530 | 0.28 |
|  | Gypsy/DIRS1 | 40073 | 40153419 | 3.22 |
|  | Retroviral | 0 | 0 | 0 |
| DNA transposons: | | 540865 | 92192402 | 7.39 |
|  | hobo-Activator | 217274 | 13882829 | 1.11 |
|  | Tc1-IS630-Pogo | 143356 | 50970391 | 4.09 |
|  | En-Spm | 0 | 0 | 0 |
|  | MULE-MuDR | 2093 | 567999 | 0.05 |
|  | PiggyBac | 4731 | 745444 | 0.06 |
|  | Tourist/Harbinger | 17398 | 3155716 | 0.25 |
|  | Other (Mirage, P-element, Transib) | 3833 | 1889126 | 0.15 |
| Rolling-circles: | | 697280 | 71292930 | 5.72 |
| Unclassified: | | 2612322 | 153887328 | 12.34 |
| Total interspersed repeats: | |  | 629094211 | 50.45 |
| Small RNA: | | 109245 | 8407915 | 0.67 |
| Satellites: | | 5 | 7657 | 0 |
| Simple repeats: | | 271971 | 14663156 | 1.18 |
| Low complexity: | | 32057 | 1567111 | 0.13 |

Table S4a. Metadata for 15 *Satyrium semiluna* specimens for which whole-genome sequence data were generated. Four of the Alberta individuals and all seven BC individuals were immediately placed in 95% after collection and stored at -20°C until DNA extraction. All four MT individuals were physically euthanized, dried in glassine envelopes, stored at room temperature for one month, and then stored at -20°C until extraction.

UASM IDs are unique identifiers for all specimens in the Univeristy of Alberta’s E. H. Strickland Entomological Museum. Wings are preserved in glassine envelopes and remainng soft tissues in 95% ethanol.

| Individual ID | UASM ID | Sex | Province or state | Location name |
| --- | --- | --- | --- | --- |
| semiluna_009 | UASM423534 | female | Alberta | Blakiston Flan |
| semiluna_010 | UASM423535 | male | Alberta | Blakiston Flan |
| semiluna_012 | UASM423536 | male | Alberta | Blakiston Flan |
| semiluna_016 | UASM423537 | male | Alberta | Blakiston Flan |
| semiluna_004 | UASM423538 | unsexed | British Columbia | Richter Pass |
| semiluna_005 | UASM423539 | unsexed | British Columbia | Richter Pass |
| semiluna_007 | UASM423540 | unsexed | British Columbia | Richter Pass |
| semiluna_008 | UASM423541 | unsexed | British Columbia | Richter Pass |
| semiluna_024 | UASM423542 | unsexed | British Columbia | Anarchist Mountain |
| semiluna_025 | UASM423543 | unsexed | British Columbia | Anarchist Mountain |
| semiluna_026 | UASM423544 | unsexed | British Columbia | Anarchist Mountain |
| semiluna_027 | UASM423545 | unsexed | Montana | 20 km SW of Red Lodge |
| semiluna_030 | UASM423546 | unsexed | Montana | 20 km SW of Red Lodge |
| semiluna_032 | UASM423547 | unsexed | Montana | 20 km SW of Red Lodge |
| semiluna_033 | UASM423548 | unsexed | Montana | 20 km SW of Red Lodge |

Table S4b. Continuation of Table S1a. Total reads are the cumulative number of unfiltered Illumina reads assigned to each individual.

| Individual ID | Latitude | Longitude | Collector | Collection date | Total reads |
| --- | --- | --- | --- | --- | --- |
| semiluna_009 | 49.079 | -113.866 | James Glasier | 14-Jul-21 | 71500620 |
| semiluna_010 | 49.078 | -113.869 | James Glasier | 14-Jul-21 | 79637697 |
| semiluna_012 | 49.076 | -113.869 | James Glasier | 14-Jul-21 | 83669611 |
| semiluna_016 | 49.068 | -113.877 | James Glasier | 14-Jul-21 | 85483727 |
| semiluna_004 | 49.085 | -119.600 | James Glasier | 21-Jun-21 | 102303395 |
| semiluna_005 | 49.085 | -119.600 | James Glasier | 21-Jun-21 | 110863887 |
| semiluna_007 | 49.084 | -119.601 | James Glasier | 21-Jun-21 | 67866061 |
| semiluna_008 | 49.084 | -119.601 | James Glasier | 21-Jun-21 | 128053215 |
| semiluna_024 | 49.007 | -119.370 | James Glasier | 19-Jun-21 | 64952222 |
| semiluna_025 | 49.007 | -119.409 | James Glasier | 19-Jun-21 | 98258363 |
| semiluna_026 | 49.007 | -119.369 | James Glasier | 19-Jun-21 | 87786345 |
| semiluna_027 | 45.054 | -109.422 | Steve Kohler | 13-Jun-21 | 95634877 |
| semiluna_030 | 45.054 | -109.422 | Steve Kohler | 13-Jun-21 | 76413308 |
| semiluna_032 | 45.054 | -109.422 | Steve Kohler | 13-Jun-21 | 74551610 |
| semiluna_033 | 45.054 | -109.422 | Steve Kohler | 13-Jun-21 | 81529947 |

Table S5. The relative contributions of each variable to *Satyrium semiluna* niche models, estimated as mean permutational importance based on AUC values and mean percent contribution.

| Variable | Permutation importance | Percent contribution |
| --- | --- | --- |
| mean summer precipitation | 41.1 | 29.9 |
| mean autumn precipitation | 18.9 | 8.8 |
| mean spring precipitation | 12.9 | 7.6 |
| mean spring temperature | 5.9 | 4.3 |
| mean relative humidity | 5.7 | 10.4 |
| terrain ruggedness | 4 | 11.9 |
| cooling degree days | 4 | 3.8 |
| mean annual precipitation | 3.3 | 1.1 |
| precipitation as snow | 1.9 | 5 |
| land cover | 0.9 | 2.1 |
| mean summer temperature | 0.7 | 0.6 |
| heatload | 0.4 | 0.9 |
| mean winter temperture | 0.3 | 0.2 |
| continentality | 0.2 | 0.2 |
| mean winter precipitation | 0 | 11.9 |
| number of frost free days | 0 | 0.9 |
| mean autumn temperature | 0 | 0.7 |
| growing degree days | 0 | 0 |
| mean annual temperature | 0 | 0 |

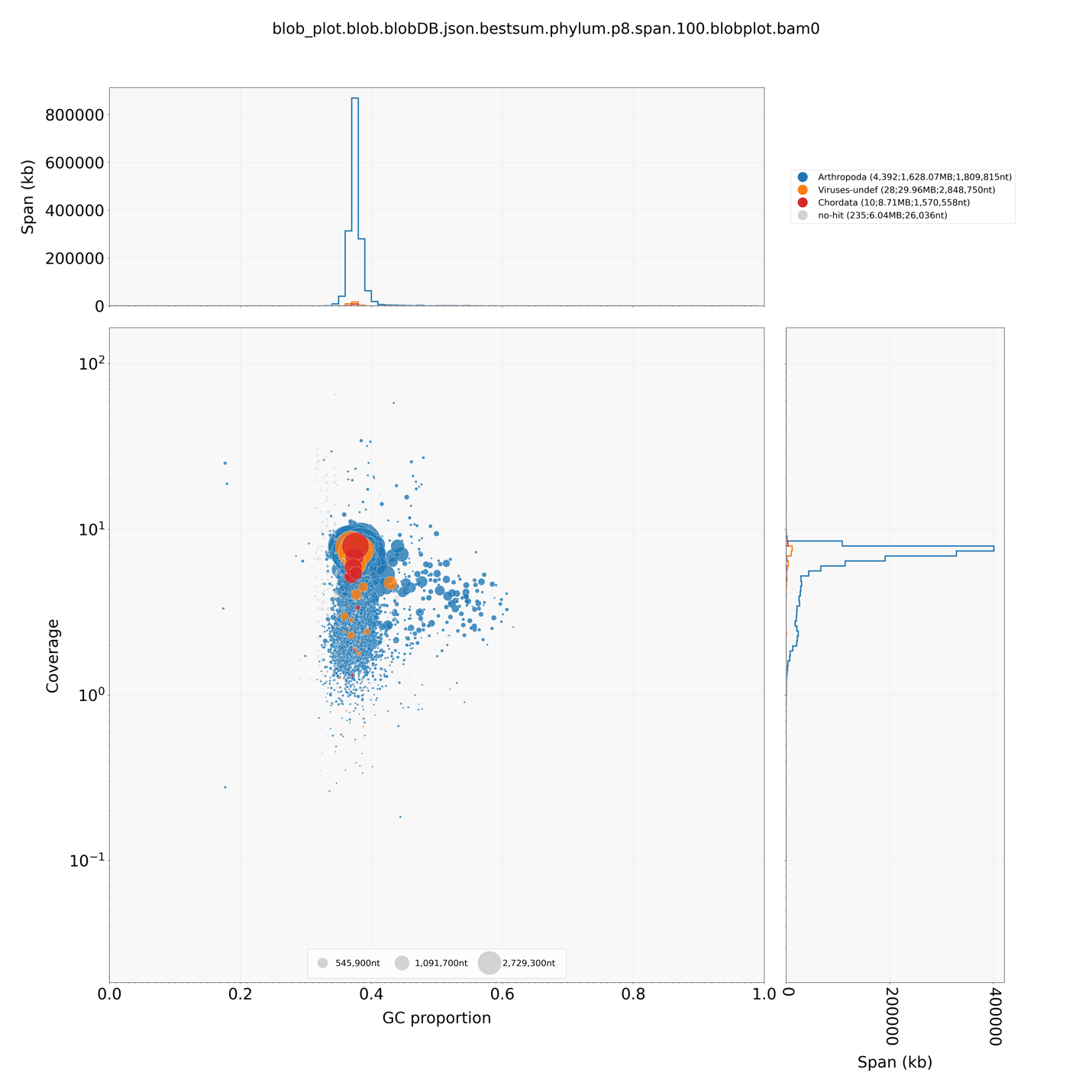

Figure S1. BlobToolkit results, showing off-target and microbial contigs.

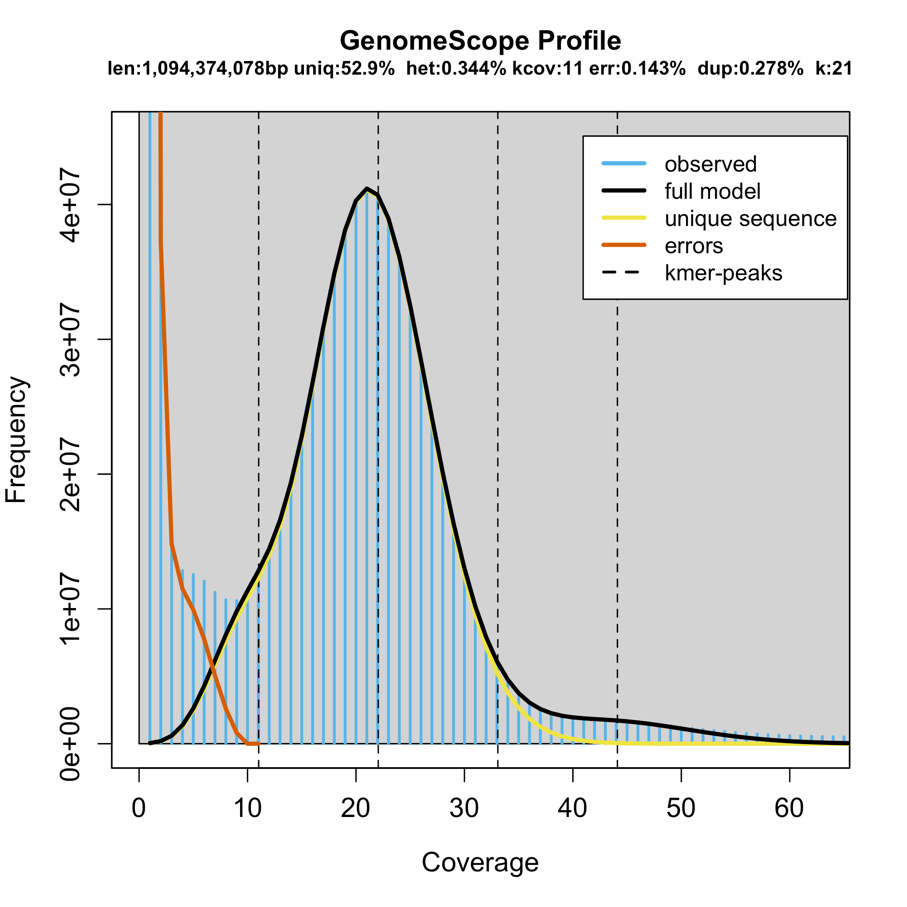

Figure S2. *k*-mer profile (*k* = 21) of the *S. semiluna* genome debarcoded raw reads as calculated by Jellyfish and GenomeScope. Blue indicates observed distribution and black, yellow, and red lines indicate modeled *k*-mer distributions of the full genome, unique sequences, and sequencing errors, respectively. Statistics for assembly length, unique sequences, heterozygosity, *k­*-mer coverage, error, and duplicated sequences provided at the top.

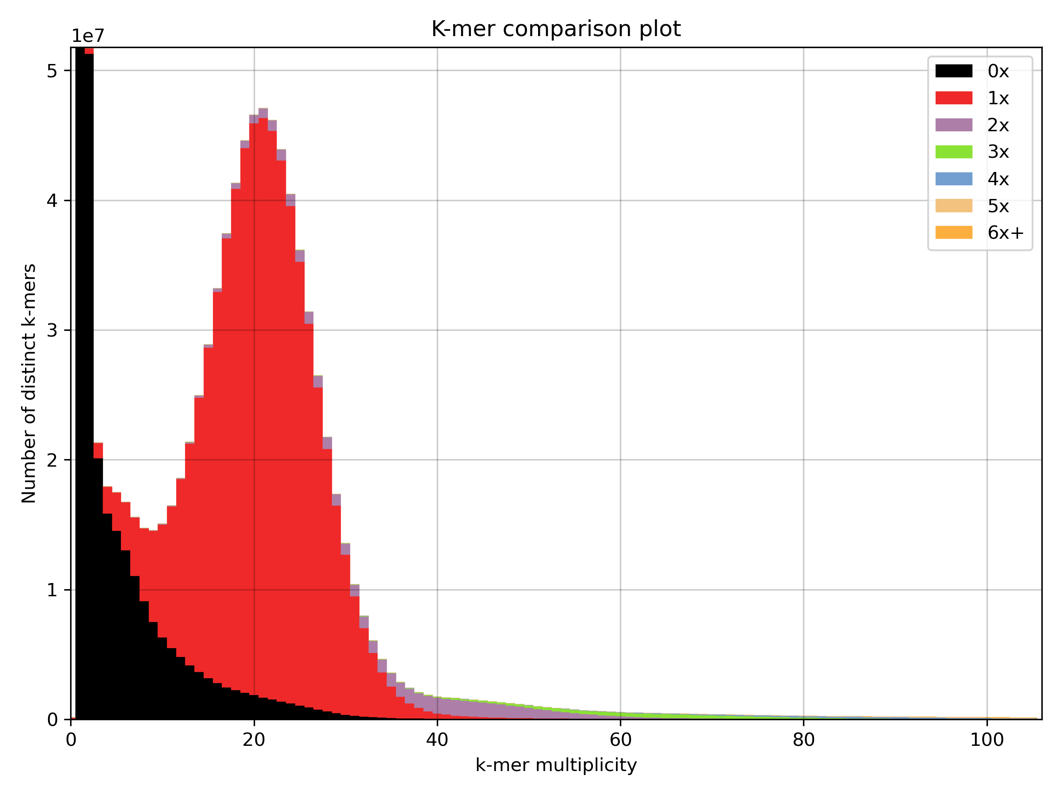

Figure S3. *k*-mer spectra comparison of raw reads to the genome assembly (color represents copy number, black indicates content not present in the assembly).

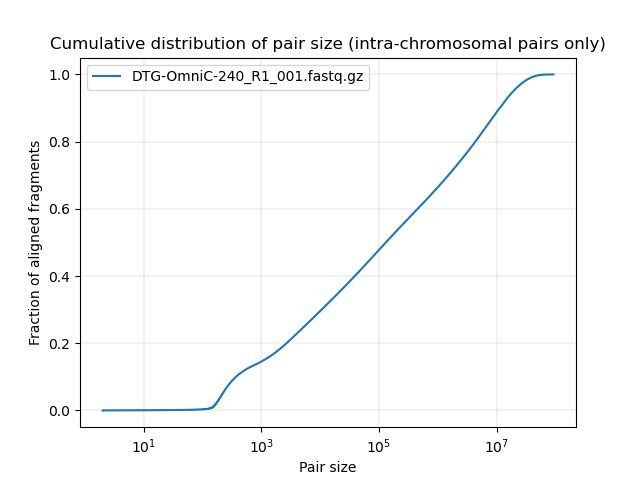

Figure S4. Cumulative distribution of intra-chromosomal pair size from HiRise.

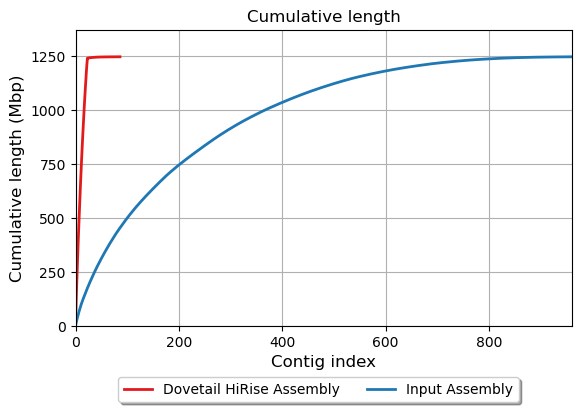

Figure S5. HiRise assembly contiguity as compared to input assembly.

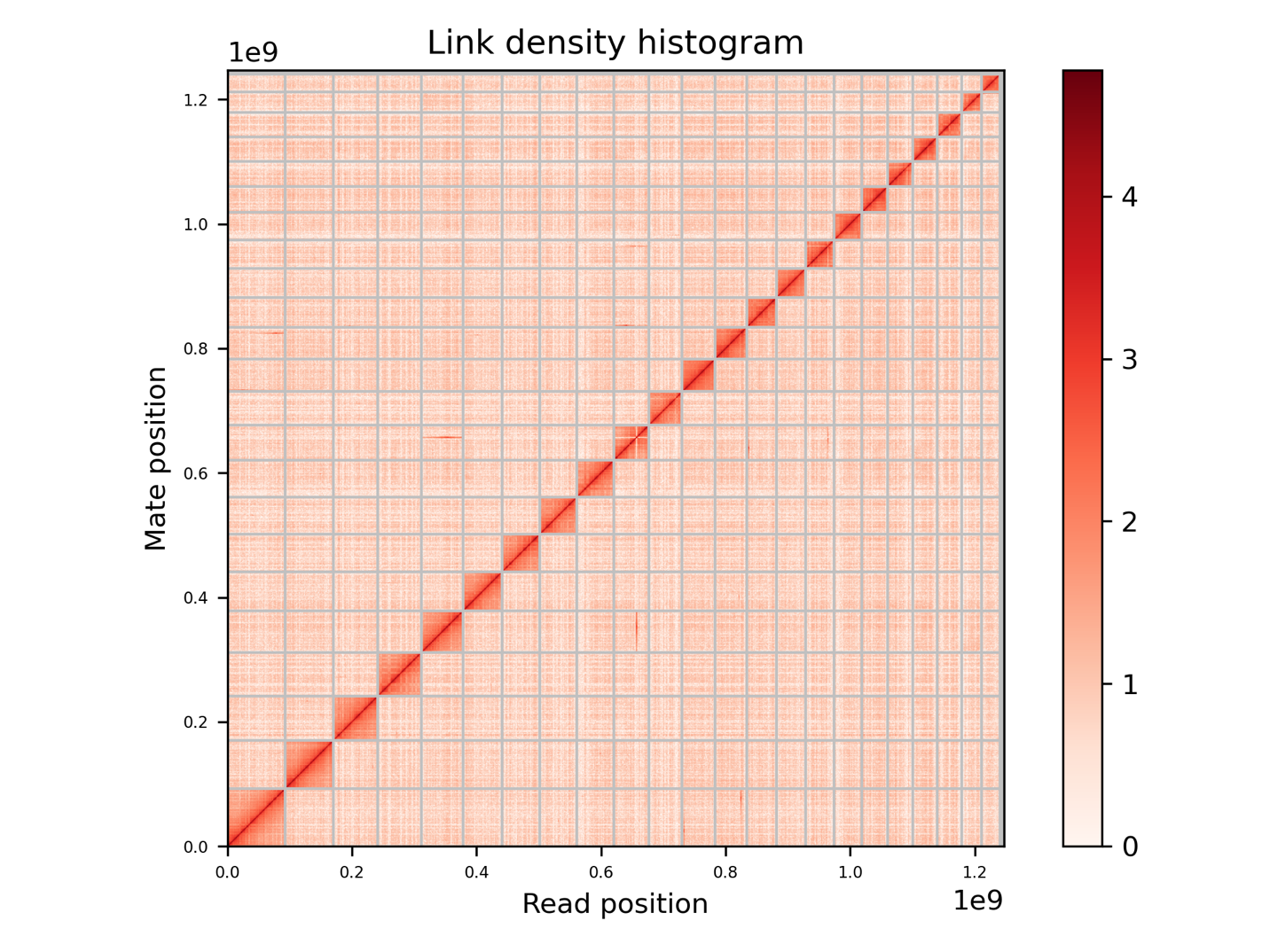

Figure S6. A Hi-C contact map for the *Satyrium semiluna* genome assembly. Each cell in the contact map corresponds to sequencing data supporting the linkage (or join) between two of such regions. Scaffolds (putative chromosomes, *n* = 21) are separated by grey lines.

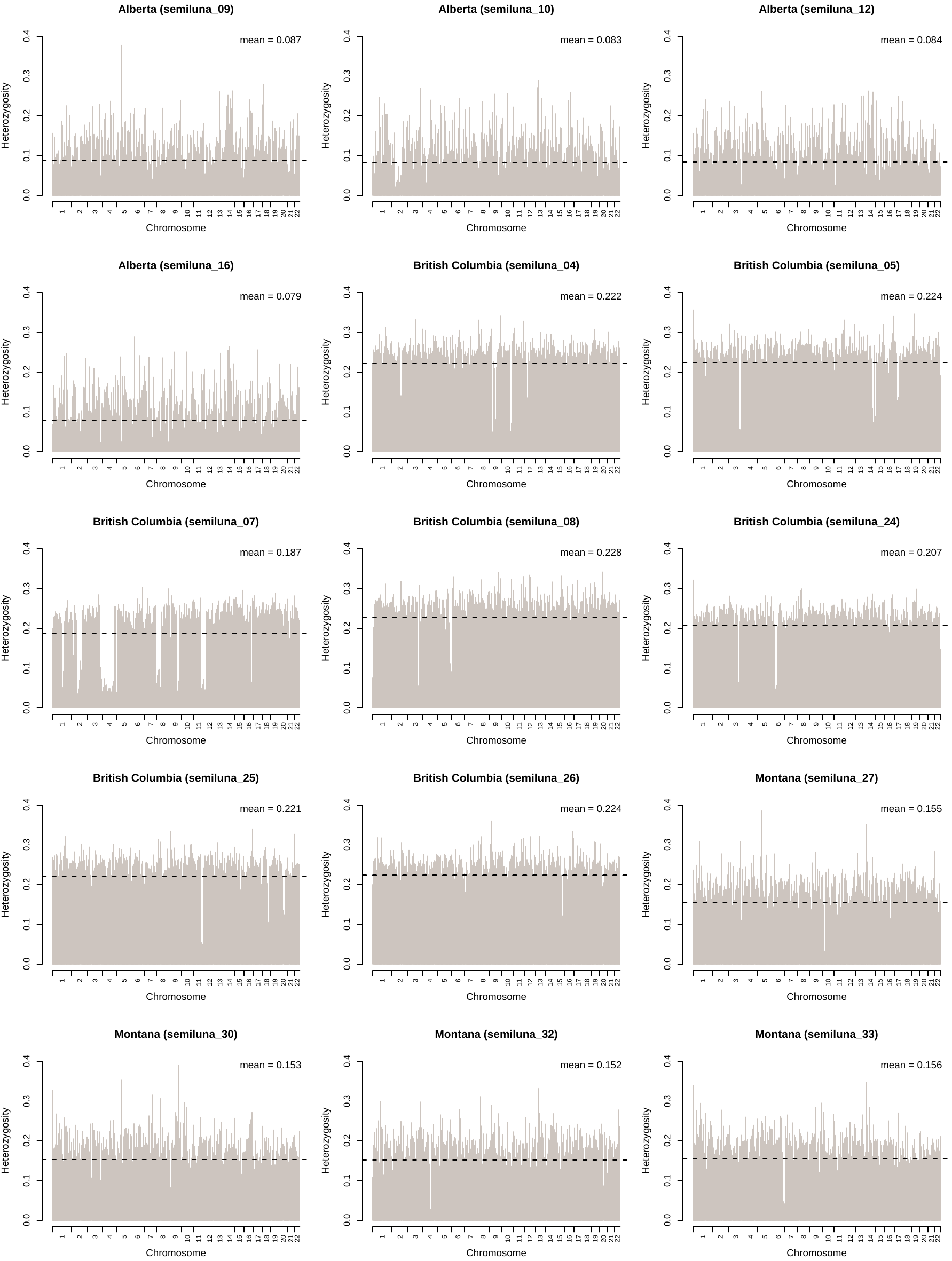

Figure S7. Heterozygosity within nonoverlapping 1 MB windows across the autosomal genome of all 15 sequenced *Satyrium semiluna* individuals. Mean genome-wide heterozygosity was estimated for each individual by averaging heterozygosity across all windows.

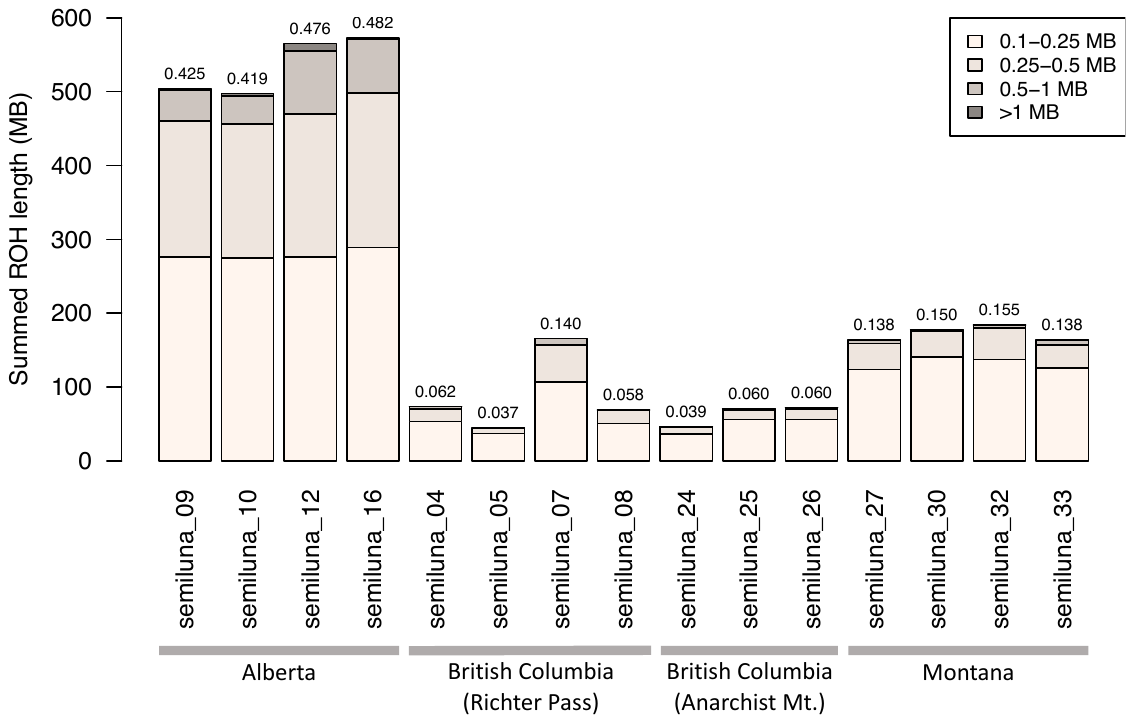

Figure S8. Cumulative length of runs of homozygosity (ROH) for each individual, categorized according to ROH length. Unlike ROH analyses presented in the main text, here we first masked all repetitive sequences identified in RepeatMasker analyses (Table S3), effectively removing them from ROH analyses, and filtered for minor allele frequencies <0.05. This renders ROH estimates more compared with those of a few other studies; e.g., addressing Channel Island foxes (Robinson et al. 2016; 2018) as well as moose (Kyriazis et al. 2023) and gray wolf populations (Kardos et al. 2018; Robison et al. 2019) on Isle Royale. An estimate of inbreeding, FROH, calculated as the total length of all ROH calls >0.1 MB divided by the cumulative length of the autosomal genome, is reported above each individual’s bar. This statistic approximates the proportion of the genome for which both copies of region are identical by descent. These analyses indicate that historical inbreeding in the Alberta *S. semiluna* population is among the highest ever recorded in a wild popualtion.
